# Semaglutide engages distinct brainstem-to-hypothalamus circuits to suppress motivated feeding and regulate ketogenesis and energy expenditure

**DOI:** 10.64898/2026.09.04.749561

**Authors:** Sebastian Blid Sköldheden, Júlia Teixidor-Deulofeu, Johan Ruud, Linda Engström Ruud

## Abstract

Semaglutide-induced weight loss requires neurons in the dorsal vagal complex (DVC), but how DVC-derived downstream brain circuits coordinate the drug’s effects on feeding and metabolism is unknown. We show that semaglutide suppresses fasting-induced AgRP neuron activation through DVC neurons, including *Adcyap1*+ neurons of the nucleus of the solitary tract (*Adcyap1*^NTS^). Projection-specific optogenetic stimulation reveals that *Adcyap1*^NTS^ inputs to the arcuate nucleus and dorsomedial hypothalamus non-aversively suppress feeding during elevated motivational drive, while sparing active-phase chow intake. *Adcyap1*^NTS^→arcuate stimulation additionally promotes food intake-independent ketogenesis and weight loss, while stimulation of the *Adcyap1*^NTS^→DMH pathway lowers energy expenditure. Crucially, stimulation of semaglutide-responsive NTS projections to both hypothalamic regions recapitulates key effects of the corresponding *Adcyap1*^NTS^ pathways on palatable-food intake and metabolism, while also suppressing fasting-induced AgRP neuron activation, demonstrating that these pathway-specific functions are retained within neuronal circuits recruited by semaglutide. Together, our findings identify brainstem-to-hypothalamus circuit substrates through which semaglutide regulates motivated feeding and metabolic state downstream of the DVC.

## INTRODUCTION

Semaglutide, a long-acting glucagon-like peptide-1 (GLP-1) receptor agonist (GLP-1RA), has transformed the pharmacological management of overweight and obesity and lowers body weight primarily through centrally mediated mechanisms^1–3^. Recent work has demonstrated hypothalamic uptake of long-acting GLP-1RAs^4,5^ and identified hypothalamic GLP-1R-expressing neuronal populations that contribute to their anorectic effects, such as thyrotropin-releasing hormone (TRH)-expressing neurons in the arcuate nucleus (ARC), which suppress the activity of orexigenic AgRP neurons^6^. In addition, in the dorsomedial hypothalamus (DMH), GLP-1R-expressing inhibitory neurons are activated by GLP-1R agonism, inhibit AgRP neuronal activity and feeding behavior^7^, and are sufficient to suppress feeding in response to GLP-1R agonism^8^. Thus, in the context of GLP-1R agonism, the ARC and DMH populations identified to date ultimately converge functionally on AgRP neurons^6,7^, consistent with *in vivo* evidence that GLP-1R agonism rapidly suppresses AgRP neuron activity^9^. However, whether these local hypothalamic circuits are recruited solely through local GLP-1R signaling, or whether they can also be engaged by upstream GLP-1RA-responsive inputs from other brain regions, remains unresolved.

In parallel, the dorsal vagal complex (DVC) has emerged as a critical brainstem entry-point through which long-acting GLP-1RAs exert their weight-lowering effects^10–13^. Within this region, adenylate cyclase activating polypeptide 1 (*Adcyap1*)-expressing neurons of the nucleus of the solitary tract (NTS; *Adcyap1*^NTS^), activated by GLP-1R-expressing neurons in the area postrema (AP), mediate the drug’s weight-loss effect^11^. *Adcyap1*^NTS^ neurons further project to multiple brainstem and forebrain structures regulating food intake and metabolism^11^. However, how these distinct DVC-derived projections contribute mechanistically to the effects of semaglutide remains unknown. This question is particularly relevant given that semaglutide-responsive DVC neurons project directly to both the ARC and DMH^11^, providing an anatomical substrate through which brainstem GLP-1RA signaling could engage hypothalamic circuits. Determining the functional roles of these semaglutide-responsive projections, whether they converge functionally on AgRP neurons, and whether they carry distinct behavioral and metabolic components of drug action is critical for understanding how DVC and hypothalamic nodes are integrated in the semaglutide response.

In the present study, we hypothesized that semaglutide engages DVC neurons to suppress AgRP neuron activity, thereby influencing feeding behavior and whole-body metabolism. We find that *Adcyap1*^AP/NTS^ neurons are required for semaglutide to suppress fasting-induced activation of AgRP neurons. Consistent with a functional contribution of AgRP neuron suppression to semaglutide-induced weight loss, chemogenetic activation of AgRP neurons during subchronic semaglutide treatment in diet-induced obese (DIO) mice reversed the drug-induced reductions in food intake and body weight. We further demonstrate that stimulation of *Adcyap1*^NTS^-derived projections to the ARC or DMH selectively suppresses feeding under conditions of elevated motivational drive, and that activation of the *Adcyap1*^NTS^→ARC pathway promotes ketogenesis and weight loss independently of food intake, while activation of the *Adcyap1*^NTS^→DMH pathway instead lowers energy expenditure. Both projections also suppressed fasting-induced activation of AgRP neurons. Crucially, key effects were recapitulated by projection-specific optogenetic stimulation of semaglutide-responsive NTS neurons, thereby linking semaglutide-associated functional outputs to defined NTS→ARC and NTS→DMH pathways. We conclude that semaglutide engages functionally distinct NTS→ARC and NTS→DMH pathways capable of exerting context-dependent effects on motivated feeding, metabolic state and fasting-induced activation of AgRP neurons.

## RESULTS

### Semaglutide suppresses hunger-evoked AgRP neuron activation, and counteracting this suppression restores feeding and reverses weight loss

To study the ability of semaglutide to inhibit AgRP neurons, we paired semaglutide treatment with two stimuli known to activate AgRP neurons: systemic ghrelin administration during the inactive (light) phase when mice normally do not engage in feeding^14^ or a brief fasting period extending into the dark phase, during which mice normally eat^15^. Ghrelin markedly increased food intake, an effect that was completely abolished by prior semaglutide treatment (Figure 1A). Correspondingly, ghrelin-induced c-Fos protein expression (a marker for neuronal activation) in the ARC was substantially reduced in semaglutide-treated animals (Figures 1B and 1C), with a decrease in activated AgRP neurons as revealed by fluorescent *in situ* hybridization (Figures 1D and 1E). Thus, semaglutide counteracts the potent feeding effect of the hunger hormone ghrelin and, based on *Fos* mRNA expression, reduces ghrelin-induced activation of AgRP neurons. Furthermore, as expected, a short fast into the dark phase (Figure 1F) strongly induced c-Fos protein expression in the ARC of vehicle-treated mice, but this response was blunted by semaglutide pretreatment (Figures 1G and 1H). *In situ* hybridization confirmed reduced *Fos* mRNA expression in *Agrp* mRNA-expressing neurons (Figures 1I and 1J). These findings indicate that reduced c-Fos expression at both the protein and mRNA level provides a read-out of semaglutide-mediated suppression of AgRP neurons.

**Figure 1.**
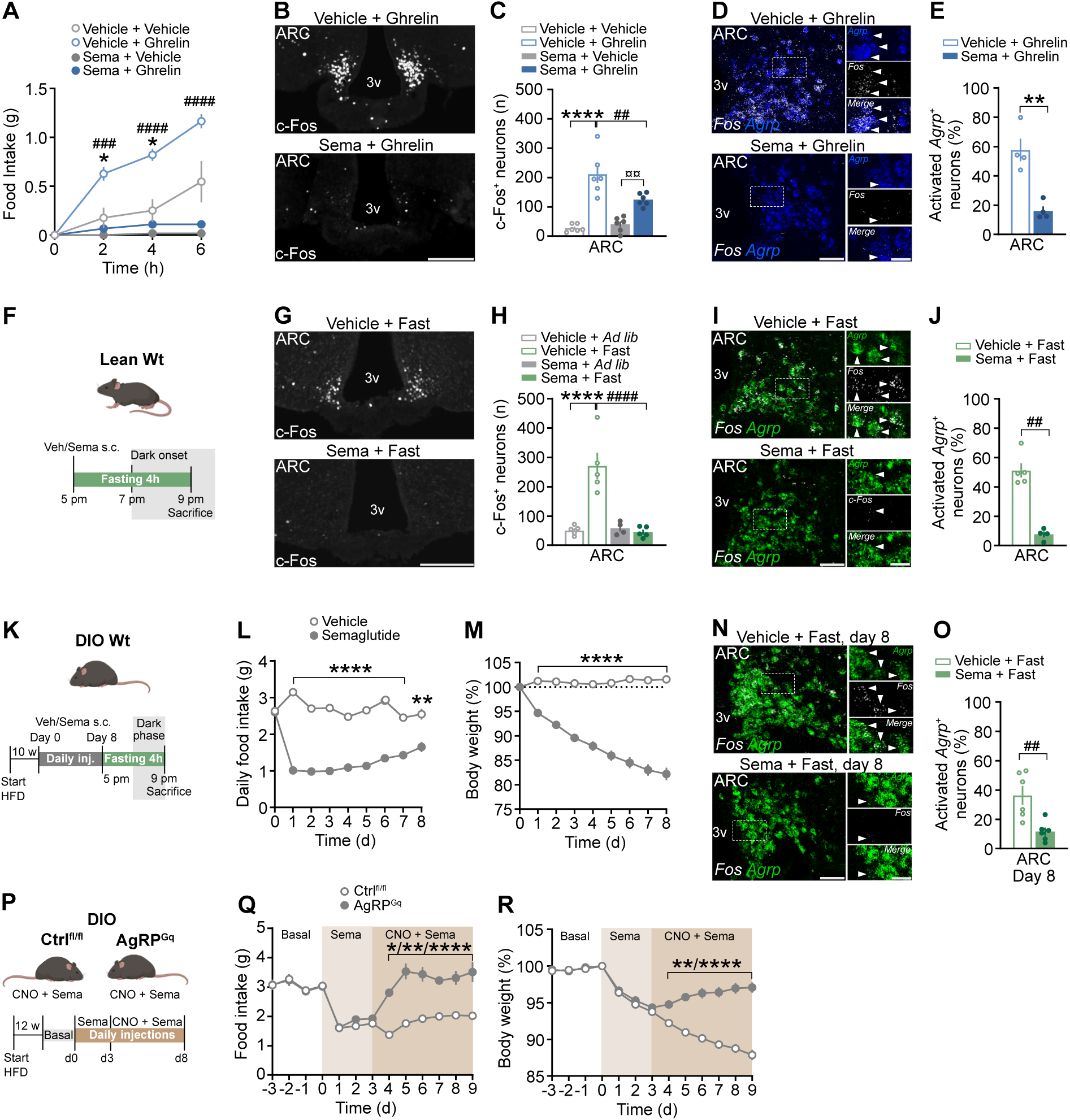
Semaglutide suppresses hunger-evoked AgRP neuron activation, and counteracting this suppression restores feeding and reverses weight loss. (A) Semaglutide effects on ghrelin-induced food intake in wt mice. *n=6*. (B) Ghrelin-induced c-Fos expression (white) in the ARC of wt mice pre-treated with vehicle or semaglutide. (C) Quantification of c-Fos expression in (B). *n=6*. (D) Fluorescent *in situ* hybridization showing *Agrp* (blue) and *Fos* (white) mRNA expression in the ARC of mice injected with ghrelin and pretreated with either vehicle or semaglutide. (E) Proportion of *Agrp*^+^ neurons co-expressing *Fos* mRNA in (D). *n=4*. (F) Approach to study the effect of semaglutide on fasting-induced activation of AgRP neurons. (G) Fasting-induced c-Fos expression (white) in the ARC of wt mice pre-treated with vehicle or semaglutide. (H) Quantification of c-Fos expression in (G) *n=4-5*. (I) Fluorescent *in situ* hybridization showing *Agrp* (green) and *Fos* (white) mRNA expression in the ARC of mice fasted into the dark phase and pretreated with either vehicle or semaglutide. (J) Proportion of *Agrp*^+^ neurons co-expressing *Fos* mRNA in (I) *n=4-5*. (K) Approach to study the effect of subchronic treatment of wt DIO mice with vehicle or semaglutide on feeding, body weight and activation of AgRP neurons. (L, M) Food intake (L) and body weight (M) in wt DIO mice treated with vehicle or semaglutide over 8 days. *n=12*. (N) Fluorescent *in situ* hybridization showing *Agrp* (green) and *Fos* (white) mRNA expression in the ARC of mice in (L, M). (O) Proportion of *Agrp*^+^ neurons co-expressing *Fos* mRNA in (N). *n=6*. (P) Approach to chemogenetically counteract semaglutide’s inhibitory effect on AgRP neurons in DIO mice (Ctrl^fl/fl^ and AgRP^Gq^). (Q, R) Food intake (Q) and body weight (R) in DIO Ctrl^fl/fl^ and AgRP^Gq^ mice treated daily with semaglutide and CNO. *n=10-15*. \**p*<0.05, **/^##^/^¤¤^*p*<0.01, ^###^*p*<0.001, ****/^####^*p*<0.0001. Data were analyzed using unpaired t-tests (E, J, O), one-way ANOVA with Šidák’s post-hoc test (C, H) or two-way ANOVA with Tukey’s (A) or Šidák’s (L, M, Q, R) post-hoc test. Error bars represent ± SEM. Boxed areas in (D, I, N) are magnified and shown next to each panel. Arrowheads point at double-labeled neurons. Scale bars = 200 µm (B, G), 50 µm (D, I, N) and 20 µm for high magnification images in (D, I, N). ARC, arcuate nucleus; DIO, diet-induced obese; 3v, third ventricle.

To determine whether this inhibitory effect is maintained under clinically relevant conditions, we next exposed DIO mice undergoing semaglutide treatment treated to the same fasting paradigm (Figure 1K). As expected, semaglutide reduced food intake and body weight over the treatment period (Figures 1L and 1M). Notably, semaglutide strongly suppressed fasting-induced AgRP neuron activation also in this setting (Figures 1N and 1O), suggesting that semaglutide-mediated suppression of AgRP neurons persists even in obese mice under prolonged exposure to the drug.

We next asked whether semaglutide-induced inhibition of AgRP neurons is required for its anorectic and weight-lowering effects. To test this, we used DIO mice, in which AgRP neurons can be chemogenetically activated through excitatory hM3Dq (AgRP^Gq^). Following daily semaglutide injections, mice received CNO at dark onset to counteract AgRP neuron inhibition (Figure 1P). While CNO-injected Ctrl^fl/fl^ mice continued to lose weight, chemogenetic activation of AgRP neurons in AgRP^Gq^ mice completely reversed semaglutide-induced reductions in food intake and changed the body weight trajectory towards weight gain (Figures 1Q and 1R). Together, these findings suggest that semaglutide suppresses hunger-promoting AgRP neurons in both acute lean and obese conditions, and that suppression of AgRP neurons is an important mediator of semaglutide-induced weight loss under the present experimental conditions.

### Semaglutide suppresses AgRP neurons through *Adcyap1*^AP/NTS^ neurons

Semaglutide acts on the DVC to reduce food intake and body weight^10–12^, and we previously found that semaglutide-activated DVC neurons project directly to hypothalamic sites involved in regulating energy balance, including the ARC and the DMH^11^. Because the DMH is known to regulate AgRP neurons^7,16,17^, and because chemogenetic reactivation of semaglutide-responsive DVC neurons activates the DMH but not the ARC^11^, we hypothesized that DVC neurons contribute to semaglutide-induced inhibition of AgRP neurons, either through direct NTS→ARC projections or indirectly via the DMH.

To test whether semaglutide-responsive DVC neurons can inhibit AgRP neurons, we first used mice in which Cre recombinase is expressed under control of the *Fos* promoter (TRAP2), allowing neurons activated by semaglutide to be permanently tagged and later manipulated. We stereotaxically injected a Cre-dependent AAV vector encoding the excitatory hM3Dq and mCherry into the DVC of TRAP2 mice (Figure 2A). Following recovery, mice received either vehicle or semaglutide, and four hours later 4-hydroxytamoxifen (4TM), which permits nuclear translocation of Cre and recombination of the AAV vector during a restricted time window. This generated two groups: vehTRAP-Gq^DVC^ (control) and semaTRAP-Gq^DVC^ mice (expressing hM3Dq specifically in semaglutide-responsive DVC neurons). This approach allowed for later chemogenetic reactivation of neurons previously engaged by semaglutide. Importantly, in vehTRAP-Gq^DVC^ mice, mCherry labeling and CNO-induced c-Fos expression in the DVC were sparse, reflecting low background recombination (Figures S1A and S1B). In contrast, semaTRAP-Gq^DVC^ mice showed robust mCherry expression and widespread CNO-induced c-Fos expression throughout all DVC subregions (Figures S1A and S1B), consistent with efficient chemogenetic reactivation of semaglutide-responsive neurons and recapitulation of the activation pattern produced by semaglutide itself^11,18^.

**Figure 2.**
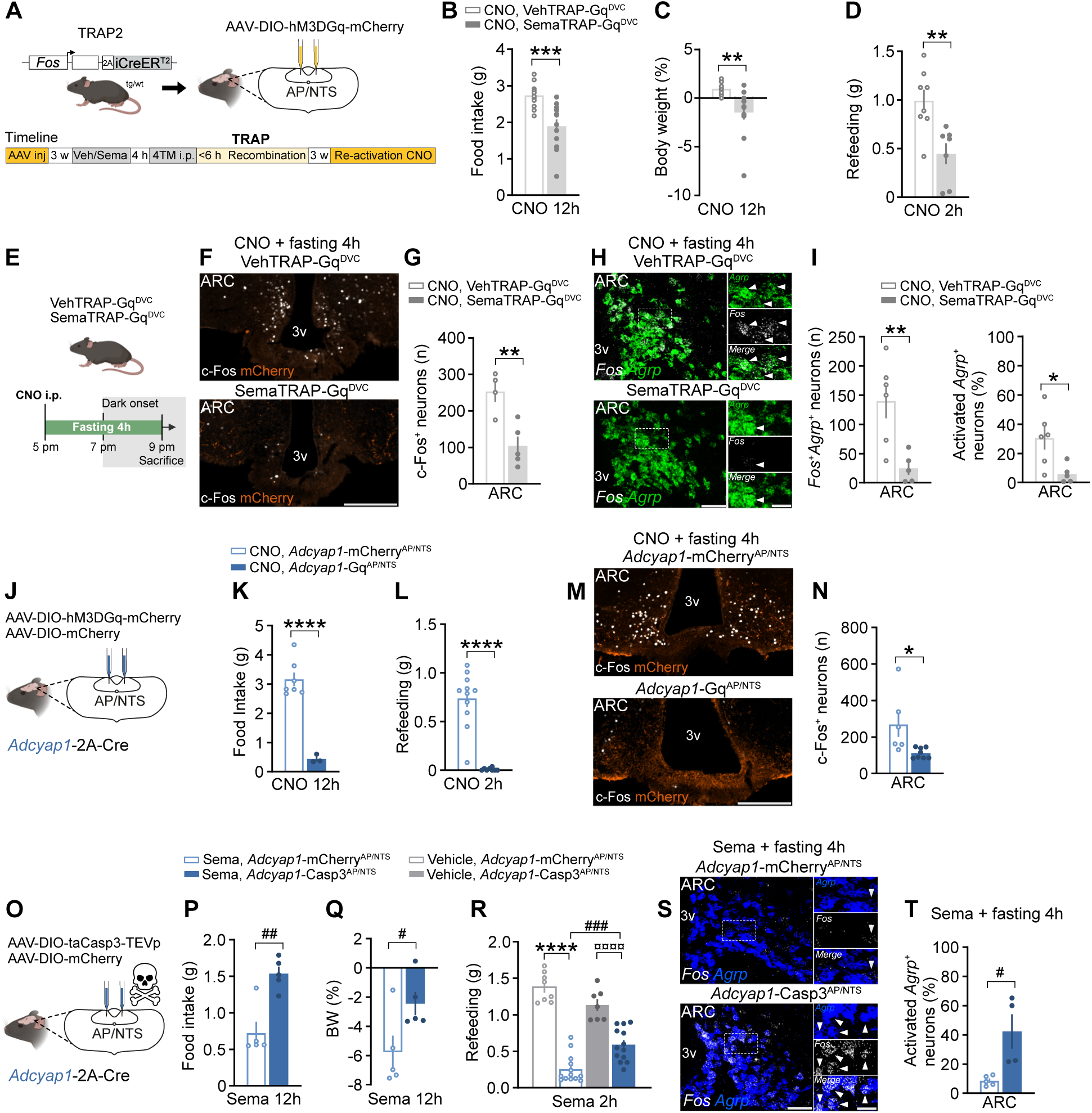
Semaglutide suppresses AgRP neurons through *Adcyap1*^AP/NTS^ neurons. (A) TRAP2-based approach to target semaglutide-responsive DVC neurons. (B, C), 12 h food intake (B) and body weight change (C) in vehTRAP-Gq^DVC^ and semaTRAP-Gq^DVC^ mice injected with CNO prior to dark onset. *n=12-14*. (D) 2 h of re-feeding in CNO-injected vehTRAP-Gq^DVC^ and semaTRAP-Gq^DVC^ mice after an overnight fast. *n=7-8*. (E) Approach to study the effect of semaglutide-responsive DVC neurons on fasting-induced activation of AgRP neurons. (F) Fasting-induced c-Fos (white) and AAV-encoded mCherry expression (orange) in the ARC of CNO-injected vehTRAP-Gq^DVC^ and semaTRAP-Gq^DVC^ mice. (G) Quantification of c-Fos expression in (F). *n=4-5*. (H) Fluorescent *in situ* hybridization showing *Agrp* (green) and *Fos* (white) mRNA expression in the ARC of vehTRAP-Gq^DVC^ and semaTRAP-Gq^DVC^ mice injected with CNO and fasted into the dark phase. (I) Quantification of *Agrp*^+^ neurons co-expressing *Fos* mRNA in (H). *n=5-6*. (J) Approach to install excitatory DREADDs in *Adcyap1*^AP/NTS^ neurons. (K, L) 12 h food intake (K) and 2 h of refeeding after an overnight fast (L) in *Adcyap1*-mCherry^AP/NTS^ and *Adcyap1*-Gq^AP/NTS^ mice after CNO injection. *n=3-7* (K) and *n=8-11* (L) (M) Fasting-induced c-Fos (white) and mCherry (orange) expression in *Adcyap1*-mCherry^AP/NTS^ and *Adcyap1*-Gq^AP/NTS^ mice after CNO injection. (N) Quantification of c-Fos expression in (M). *n=6-8*. (O) Approach to Cre-dependently delete *Adcyap1*^AP/NTS^ neurons. (P, Q) 12 h food intake (P) and body weight change (Q) in *Adcyap1*-mCherry^AP/NTS^ and *Adcyap1*-Casp3^AP/NTS^ mice treated with semaglutide prior to dark onset. *n=5*. (R) 2 h of refeeding after an overnight fast in *Adcyap1*-mCherry^AP/NTS^ and *Adcyap1*-Casp3^AP/NTS^ mice treated with vehicle or semaglutide prior to reintroduction of food. *n=7-13*. ^(S)^ Fluorescent *in situ* hybridization showing *Agrp* (blue) and *Fos* (white) mRNA expression in the ARC of *Adcyap1*-mCherry^AP/NTS^ and *Adcyap1*-Casp3^AP/NTS^ injected with semaglutide and fasted into the dark phase. (T) Proportion of *Agrp*^+^ neurons co-expressing *Fos* mRNA in (S). *n=4-5*. *^/#^*p*<0.05, **^/##^*p*<0.01, ***^/###^*p*<0.001, ****^/¤¤¤¤^*p*<0.0001. Data were analyzed using unpaired t-tests (B-D, G, I, K, L, N, P, Q, T) or one-way ANOVA with Šidák’s post-hoc test (R). Error bars represent ± SEM. Boxed areas in (H, S) are magnified and shown next to each panel. Scale bars = 200 µm (F, M), 50 µm (H, S) and 20 µm for high magnification images in (H, S). Arrowheads point at double-labeled neurons. AP, area postrema; ARC, arcuate nucleus; NTS, nucleus of the solitary tract; TRAP, targeted recombination in activated populations; 3v, third ventricle. Relates to Figure S1.

Chemogenetic stimulation of semaglutide-responsive DVC neurons reduced dark-phase feeding and body weight (Figures 2B and 2C) and, notably, suppressed fasting-induced refeeding, a strongly AgRP neuron-driven behavior^19^ (Figure 2D). This manipulation also reduced fasting-induced c-Fos expression in the ARC (Figures 2E-2G), including a marked decrease in activated AgRP neurons, in contrast to CNO-injected vehTRAP-Gq^DVC^ mice that showed robust *Fos* expression in AgRP neurons following fasting (Figures 2H and 2I). These results demonstrate that chemogenetic reactivation of semaglutide-responsive DVC neurons alone is sufficient to mimic semaglutide-induced suppression of fasting-induced AgRP neuron activity and to blunt highly motivated feeding driven by a negative energetic state.

We previously reported that a major subset of the semaglutide-activated NTS neurons express *Adcyap1* mRNA and that these *Adcyap1*^NTS^ neurons, together with a much smaller population of *Adcyap1*^+^ neurons in the AP, are necessary for the full weight-lowering effect of semaglutide in both lean and DIO mice^11^. We therefore tested if *Adcyap1*^AP/NTS^ neurons contribute to semaglutide-induced inhibition of AgRP neurons. *Adcyap1*-2A-Cre mice were stereotactically injected into the DVC with an AAV5-hSyn-DIO-hM3DGq-mCherry or the corresponding control virus encoding mCherry only (Figure 2J). Chemogenetic activation of the broad *Adcyap1*^AP/NTS^ neuronal population, which also include neurons non-responsive to semaglutide, induced c-Fos in the AP and NTS (Figures S1C and S1D) and robustly suppressed dark-phase feeding (Figure 2K), and completely blocked fasting-induced refeeding (Figure 2L). This was accompanied by a significant reduction in fasting-induced ARC c-Fos expression (Figures 2M and 2N). These findings collectively suggest that semaglutide-responsive DVC neurons, including *Adcyap1*-expressing neurons, pose direct or indirect inhibitory control over AgRP neurons.

We next assessed whether *Adcyap1*^AP/NTS^ neurons are necessary for semaglutide-induced inhibition of AgRP neurons. To this end, we specifically deleted *Adcyap1*^AP/NTS^ neurons by injecting a Cre-dependent AAV5-EF1a-DIO-taCasp3-TEVp vector^20^ or a control vector encoding mCherry into the DVC of *Adcyap1*-2A-Cre mice (Figure 2O). In Cre-positive neurons, TEVp (Tobacco Etch Virus protease) cleaves pro-taCasp3 to yield an active Caspase 3 protein that induces apoptosis^20^. This approach has previously been validated by us in the semaglutide setting, showing that deletion of *Adcyap1*^AP/NTS^ neurons significantly reduces semaglutide-induced activation of the AP and NTS^11^, and yielded two experimental groups: *Adcyap1-*Casp3^AP/NTS^ (lacking *Adcyap1^+^* AP/NTS neurons) and *Adcyap1-* mCherry^AP/NTS^ (intact controls). The AAV5 serotype was chosen to avoid deleting the efferent vagal motor neurons of the DMV, known to partly express *Adcyap1* mRNA^21^. Proper deletion of *Adcyap1*^AP/NTS^ neurons, but intact expression of *Adcyap1* mRNA in DMV neurons, was verified post-mortem using *in situ* hybridization (Figures S1E and S1F). Loss of *Adcyap1*^AP/NTS^ neurons attenuated the feeding and weight-lowering effects of semaglutide (Figures 2P and 2Q), confirming published results^11^. Notably, semaglutide-mediated suppression of fasting-induced refeeding was significantly attenuated in ablated mice (Figure 2R). Consistent with this behavioral effect, semaglutide no longer effectively suppressed fasting-induced *Fos* expression in AgRP neurons in ablated animals (Figures 2S and 2T, S1G and S1H). Accordingly, ablated mice exhibited increased numbers and proportions of activated AgRP neurons compared to intact controls. Together, these findings establish *Adcyap1*^AP/NTS^ neurons as a necessary brainstem relay linking semaglutide to suppression of fasting-induced refeeding and AgRP neuron activation.

### Activation of ARC and DMH projections from *Adcyap1*^NTS^ neurons does not suppress dark-phase feeding

We next sought to determine the functional relevance of *Adcyap1*^AP/NTS^ mediated inhibition of AgRP neurons, and whether this effect is mediated through direct projections to the ARC or indirectly via the DMH. The NTS sends widespread ascending projections throughout the brain^22,23^, including the ARC and the DMH^11,24,25^, whereas the AP exhibits a much more restricted projection pattern, primarily targeting the NTS, parabrachial nucleus and the ventrolateral medulla^26^. In addition, *Adcyap1*^+^ neurons are far more abundant in the NTS than in the AP. Together, these anatomical considerations suggest that the *Adcyap1*-derived hypothalamic projections examined here predominantly arise from the *Adcyap1*^NTS^ population.

To dissect the functional significance of these pathways on different aspects of energy balance, *Adcyap1*-2A-Cre mice received stereotaxic injections of a Cre-dependent AAV encoding channelrhodopsin-2 (ChR2) and mCherry into the NTS, followed by unilateral implantation of a fiberoptic cannula above either the ARC or the DMH (Figure 3A). Correct AAV expression, optic fiber placement above the ARC or DMH and the presence of *Adcyap1*^NTS^-derived axonal fibers in both target regions were verified histologically (Figures S2A and S2B). Optogenetic testing was conducted using a counterbalanced laser ON/OFF paradigm.

**Figure 3.**
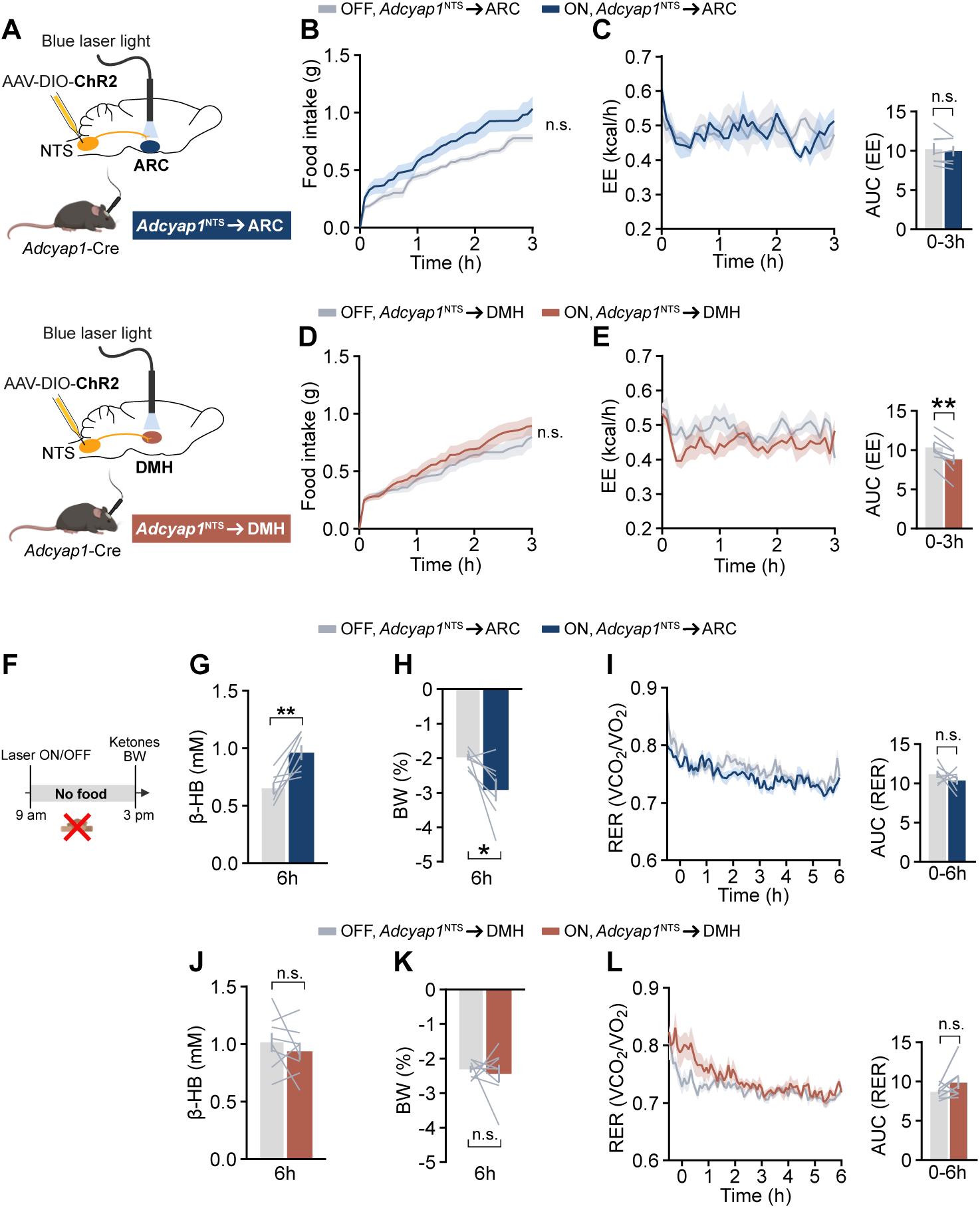
Functionally specialized *Adcyap1*^NTS^→ARC and *Adcyap1*^NTS^→DMH projections differentially regulate metabolism without altering baseline dark-phase feeding. (A) Approach to optogenetically target *Adcyap1*^NTS^ projections to either the ARC (blue) or the DMH (orange). (B, C) Dark phase food intake (B), and energy expenditure (C) in photostimulated and non-stimulated *Adcyap1*^NTS^→ARC mice. *n=7*. (D, E) Dark phase food intake (D) and energy expenditure (E) in photostimulated and non-stimulated *Adcyap1*^NTS^→DMH mice. *n=8*. (F) Approach to study the effect of photostimulation on ketogenesis and body weight in the absence of caloric intake during the light phase. (G-I) Effect of light phase photostimulation on circulating β-hydroxybutyrate levels (G), body weight change (H) and respiratory exchange ratio (I) in *Adcyap1*^NTS^→ARC mice. *n=7*. (J-L) Effect of light phase photostimulation on circulating β-hydroxybutyrate levels (J), body weight change (K) and respiratory exchange ratio (L) in *Adcyap1*^NTS^→DMH mice. *n=8*. n.s.=not significant, \**p*<0.05, \*\**p*<0.01. Data were analyzed using paired t-test (C, E, G-L) or two-way ANOVA with Šidák’s post-hoc test (B, D). Error bars represent ± SEM. ARC, arcuate nucleus; DMH, dorsomedial hypothalamus; NTS, nucleus of the solitary tract. Relates to Figure S2.

We first assessed how stimulation of *Adcyap1*^NTS^→ARC and *Adcyap1*^NTS^→DMH projections affect normal dark-phase feeding, respiratory exchange ratio (RER) and energy expenditure (EE). For this purpose, mice were housed in an open-cage indirect calorimetry system equipped with FED3.1 devices for automated, time-stamped retrieval of 20 mg food pellets^27^. Contrary to our hypothesis, photostimulation of the *Adcyap1*^NTS^→ARC pathway did not suppress dark-phase feeding or significantly alter EE (Figures 3B and 3C, and had no effect on RER (Figure S2C). Cumulative food intake was in fact numerically higher during laser ON compared with laser OFF, although the main effect of stimulation did not reach statistical significance (*p*=0.0741). Similarly, *Adcyap1*^NTS^→DMH stimulation did not suppress dark-phase feeding (Figure 3D) or significantly alter RER (Figure S2D); however, in contrast to the ARC pathway, it significantly reduced EE (Figure 3E). Together, these findings indicate that stimulation of *Adcyap1*^NTS^ projections to the ARC or DMH is insufficient to suppress physiological dark-phase feeding. However, the two projections exert pathway-specific effects on metabolic output, with the *Adcyap1*^NTS^→DMH projection selectively lowering energy expenditure independently of changes in food intake. This reveals a functional dissociation between the effects of *Adcyap1*^NTS^-derived projections on baseline feeding and metabolic output.

### The *Adcyap1*^NTS^→ARC, but not *Adcyap1*^NTS^→DMH, pathway drives ketogenesis and weight loss, promoting a fasting-like metabolic state

Given the observed dissociation between feeding and energy expenditure, we next asked whether *Adcyap1*^NTS^ projections regulate fuel utilization and energy expenditure in the absence of feeding. While optogenetic stimulation did not alter RER in free-feeding animals, ongoing nutrient intake may mask changes in the utilization of endogenous energy stores. Because AgRP neurons regulate peripheral substrate utilization in addition to feeding^28^, we asked whether stimulation of *Adcyap1*^NTS^ hypothalamic projections promotes a metabolic shift toward fat utilization. To test this, we optogenetically stimulated *Adcyap1*^NTS^→ARC or *Adcyap1*^NTS^→DMH projections for 6 h during the light phase in the absence of food (Figure 3F). Photostimulation of the *Adcyap1*^NTS^→ARC projection significantly increased circulating β-hydroxybutyrate levels, consistent with enhanced ketogenesis, and promoted weight loss (Figures 3G and 3H). This manipulation tended to reduce RER, although not significantly (Figure 3I), and did not alter EE (Figure S2E). In contrast, stimulation of the *Adcyap1*^NTS^→DMH pathway had no effect on ketogenesis, body weight, RER or EE (Figures 3J-L and S2F).

Together, these findings reveal a functional specialization of *Adcyap1*^NTS^-derived hypothalamic projections. Whereas stimulation of the *Adcyap1*^NTS^→ARC pathway promotes ketogenesis and body weight loss independent of feeding, stimulation of the *Adcyap1*^NTS^◊DMH pathway selectively reduces energy expenditure under free-feeding conditions.

### Stimulation of *Adcyap1*^NTS^**→**ARC and DMH projections suppresses fasting-induced refeeding and chocolate binge-eating in a non-aversive manner

Given that stimulation of *Adcyap1*^NTS^ projections did not affect basal dark-phase feeding, we next asked whether these pathways instead regulate feeding under conditions of elevated motivational drive to eat. We therefore first examined their impact on fasting-induced refeeding, a robust behavioral response driven by negative energy balance. Indeed, optogenetic stimulation of either *Adcyap1*^NTS^→ARC or *Adcyap1*^NTS^→DMH projections during refeeding after an overnight fast significantly reduced food intake and blunted recovery of body weight (Figures 4A-D), indicating that *Adcyap1*^NTS^ -derived hypothalamic projections suppress feeding when the drive to eat is elevated due to energy deficit.

**Figure 4.**
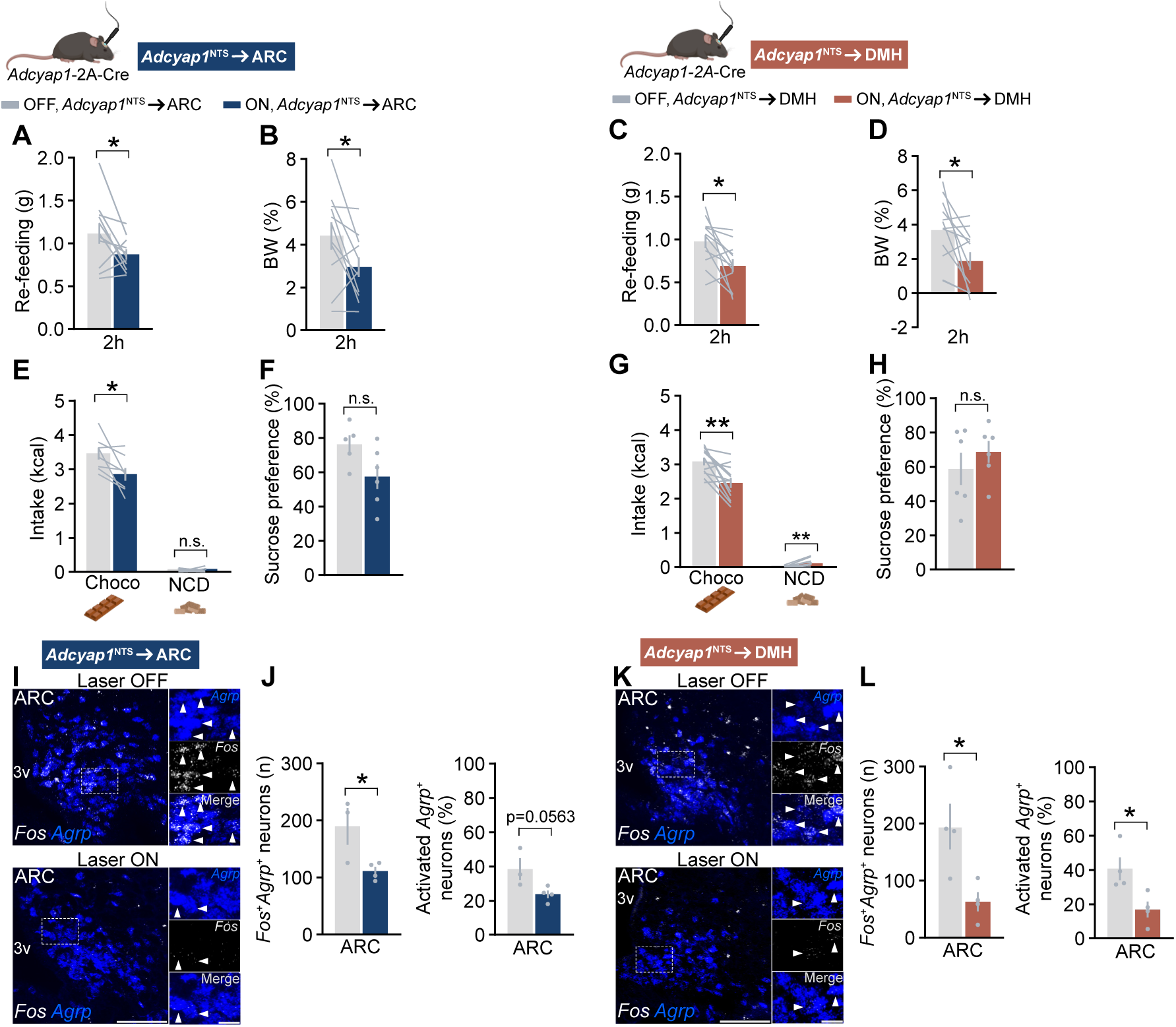
Stimulation of *Adcyap1*^NTS^→ARC and *Adcyap1*^NTS^→DMH projections suppresses motivated feeding and AgRP neuron activity in an non-aversive manner. (A, B) Effect of photostimulation on re-feeding (A) and body weight change (B) after an overnight fast in *Adcyap1*^NTS^→ARC mice. *n=11*. (C, D) Effect of photostimulation on re-feeding (C) and body weight change (D) after an overnight fast in *Adcyap1*^NTS^DMH mice. *n=12*. (E) Effect of photostimulation on binge-eating of chocolate and concomitant intake of normal chow during a 2 h restricted access paradigm in *Adcyap1*^NTS^→ARC mice. *n=8*. (F) Sucrose preference after pairing sucrose intake with photostimulation in a conditioned taste aversion protocol in *Adcyap1*^NTS^→ARC mice. *n=5-6*. (G) Effect of photostimulation of *Adcyap1*^NTS^→DMH mice on 2h chocolate and normal chow intake in the same paradigm as in (E). *n=12*. (H) Sucrose preference after pairing sucrose intake with photostimulation in a conditioned taste aversion protocol in *Adcyap1*^NTS^→DMH mice. *n=6*. (I) Fluorescent *in situ* hybridization showing *Agrp* (blue) and *Fos* (white) mRNA expression in the ARC of *Adcyap1*^NTS^→ARC mice fasted for 4 h into the dark phase with or without concomitant photostimulation. (J) Quantification of co-expression in (I). *n=3-4*. (K) Fluorescent *in situ* hybridization showing *Agrp* (blue) and *Fos* (white) mRNA expression in the ARC of *Adcyap1*^NTS^→DMH mice fasted for 4 h into the dark phase with or without concomitant photostimulation. (L) Quantification of co-expression in (K). *n=4*. n.s.=not significant, \**p*<0.05, \*\**p*<0.01. Data were analyzed using paired t-test (A-E, G), unpaired t-test (F, H, J, L). Error bars represent ± SEM. Scale bars = 100 µm (I, K) and 20 µm in magnified boxed regions (I, K). Arrowheads point at double-labeled neurons. ARC, arcuate nucleus; DMH, dorsomedial hypothalamus; NTS, nucleus of the solitary tract; 3v, third ventricle. Relates to Figure S3.

We next tested whether these projections similarly regulate the intake of palatable energy-dense food. Freely fed mice were given daily access to chocolate for 2 h during the light phase. After approximately five days, they developed a stable binge-like intake pattern, consuming >3 kcal (∼0.5 g) from chocolate while largely ignoring chow (data not shown). On day 6, photostimulation of the *Adcyap1*^NTS^→ARC pathway significantly reduced chocolate intake compared with the laser OFF condition, without altering the concurrent (and minimal) chow intake (Figure 4E). To determine whether this reduction reflected aversion or malaise, photostimulation of the same projection was paired with consumption of a palatable sucrose solution in a conditioned taste aversion (CTA) paradigm (Figure S3A). Sucrose preference during the subsequent test session did not differ between laser ON and laser OFF conditions (Figure 4F), indicating that *Adcyap1*^NTS^→ARC stimulation did not induce an aversive response to the palatable sucrose solution.

Photostimulation of the *Adcyap1*^NTS^→DMH pathway similarly reduced chocolate intake (Figure 4G), and, in contrast to ARC stimulation, produced a small but significant increase in concurrent chow intake (Figure 4G), suggesting a partial shift from palatable to homeostatic feeding. Pairing *Adcyap1*^NTS^→DMH stimulation with sucrose consumption did not alter subsequent sucrose preference (Figure 4H), indicating that the reduced chocolate intake was not associated with aversion. Together, these findings demonstrate that engagement of both the *Adcyap1*^NTS^→ARC and the *Adcyap1*^NTS^→DMH pathways suppress motivationally driven palatable food intake without inducing aversion, while the DMH pathway additionally promotes a very modest, but detectable, increase of chow intake.

### *Adcyap1*^NTS^ neurons inhibit fasting-induced *Fos* expression in AgRP neurons via direct ARC and DMH projections

We next asked whether the behavioral effects observed during optogenetic stimulation of *Adcyap1*^NTS^→ARC or *Adcyap1*^NTS^→DMH projections are associated with inhibition of AgRP neurons. Mice were subjected to a short fast into the dark phase, with concomitant photostimulation, with fasted non-photostimulated mice serving as controls. Using *in situ* hybridization, we quantified fasting-induced activation of AgRP neurons. Photostimulation of either the *Adcyap1*^NTS^→ARC (Figures 4I and 4J) or *Adcyap1*^NTS^→DMH projections (Figures 4K and 4L) significantly reduced the number and proportion of fasting-activated AgRP neurons compared to non-stimulated fasted controls. These findings demonstrate that activation of both projections ultimately impinges on AgRP neuron inhibition.

### *Adcyap1*^NTS^ neurons include both excitatory and inhibitory subpopulations, both of which are recruited by semaglutide

*Adcyap1*^NTS^ neurons have previously been reported to be glutamatergic^21^, but whether the broader *Adcyap1*^NTS^ population includes inhibitory neurons, and which subtypes are recruited by semaglutide, remains unknown. We therefore performed *in situ* hybridization on DVC sections from semaglutide-injected mice using *Slc32a1* mRNA, encoding the vesicular GABA transporter VGAT, as a marker for inhibitory neurons (Figure 5A). Approximately 30% of all *Adcyap1*^NTS^ neurons expressed *Slc32a1* mRNA and a similar proportion of *Slc32a1*^+^ neurons expressed *Adcyap1* mRNA (Figure 5B). Semaglutide robustly increased *Fos* expression in the NTS, recruiting both *Adcyap1*^+^ and *Slc32a1*^+^ populations (Figure 5C). Semaglutide administration activated about 40% of the *Adcyap1*^NTS^ neurons, and roughly one third of the inhibitory *Slc32a1*^+^ population (Figure 5C). Notably, semaglutide significantly increased the number of triple-labelled neurons, with inhibitory neurons comprising approximately half of the semaglutide-activated *Adcyap1*^NTS^ population (Figure 5D). Together, these findings indicate that semaglutide recruits both excitatory and inhibitory *Adcyap1*^NTS^ neurons, revealing previously unrecognized cellular heterogeneity within this population. Combined with our projection-specific experiments, these data further suggest that molecularly distinct *Adcyap1*^NTS^ subpopulations may differentially contribute to hypothalamic circuit regulation.

**Figure 5.**
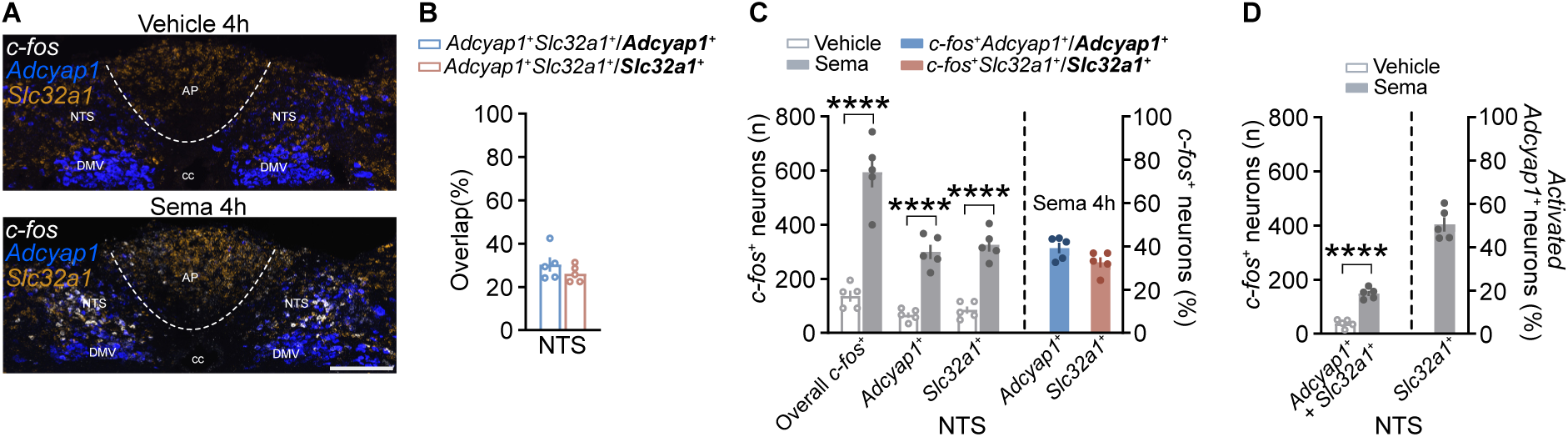
*Adcyap1*^NTS^ neurons include both excitatory and inhibitory subpopulations, both of which are recruited by semaglutide. (A) Fluorescent *in situ* hybridization showing *Adcyap1* (blue), *Slc32a1* (orange) and *Fos* (white) mRNA expression in the DVC of wt mice at 4 h after an injection with vehicle or semaglutide. (B) Quantification of the overlap between the *Adcyap1^+^* and the *Slc32a1^+^* populations in the NTS. *n=5*. (C) Quantification of semaglutide-activated neurons and their identity in terms of *Adcyap1* and *Slc32a1* mRNA expression. *n=5*. (D) Quantification of triple-labelled neurons and their relation to the activated *Adcyap1*^NTS^ population. *n=5*. \*\*\*\**p*<0.0001. Data were analyzed using unpaired t-test (C, D) with Holm-Šidák’s method to correct for multiple comparisons (C). Error bars represent ± SEM. Scale bar = 200 µm in (A). AP, area postrema; cc, central canal; DMV, dorsal motor nucleus of the vagus; NTS, nucleus of the solitary tract.

### Semaglutide-responsive NTS→hypothalamus projections are functionally specialized for motivated feeding and metabolic control

Our findings above demonstrate that *Adcyap1*^NTS^ neurons with projections to the ARC and DMH differentially regulate motivated feeding and metabolism. However, these optogenetic experiments targeted ARC and DMH projections arising from the NTS solely based on *Adcyap1* expression, which is found in a large population of neurons^11^, and therefore did not distinguish between semaglutide-responsive and semaglutide-insensitive pathways. Because semaglutide activates only a subset (∼40%) of *Adcyap1*^NTS^ neurons, we next asked whether the projection-specific functions identified above could be attributed to the semaglutide-responsive subset of neurons, and thus whether these projections constitute NTS-derived circuits through which semaglutide regulates feeding and metabolism.

To address this, we used TRAP2 mice to genetically capture semaglutide-activated NTS neurons and selectively stimulate their projections to the ARC and DMH (Figure 6A). TRAP2 mice received a Cre-dependent AAV encoding ChR2 and mCherry in the NTS, followed by optical fiber implantation unilaterally above either of the target regions (Figure 6A). Mice were then TRAPed with vehicle or semaglutide (vehTRAP^NTS^ and semaTRAP^NTS^), enabling selective expression of ChR2 in vehicle or semaglutide-activated neurons and their axonal projections for subsequent functional ChR2-assisted interrogation, during which both groups received photostimulation. Histological analysis confirmed robust TRAP-dependent AAV-mediated mCherry expression in the DVC of semaTRAP^NTS^ mice and dense terminal labeling within the targeted hypothalamic regions, while only sparser background labeling was observed in vehTRAP^NTS^ mice (Figures 6B and 6C).

**Figure 6.**
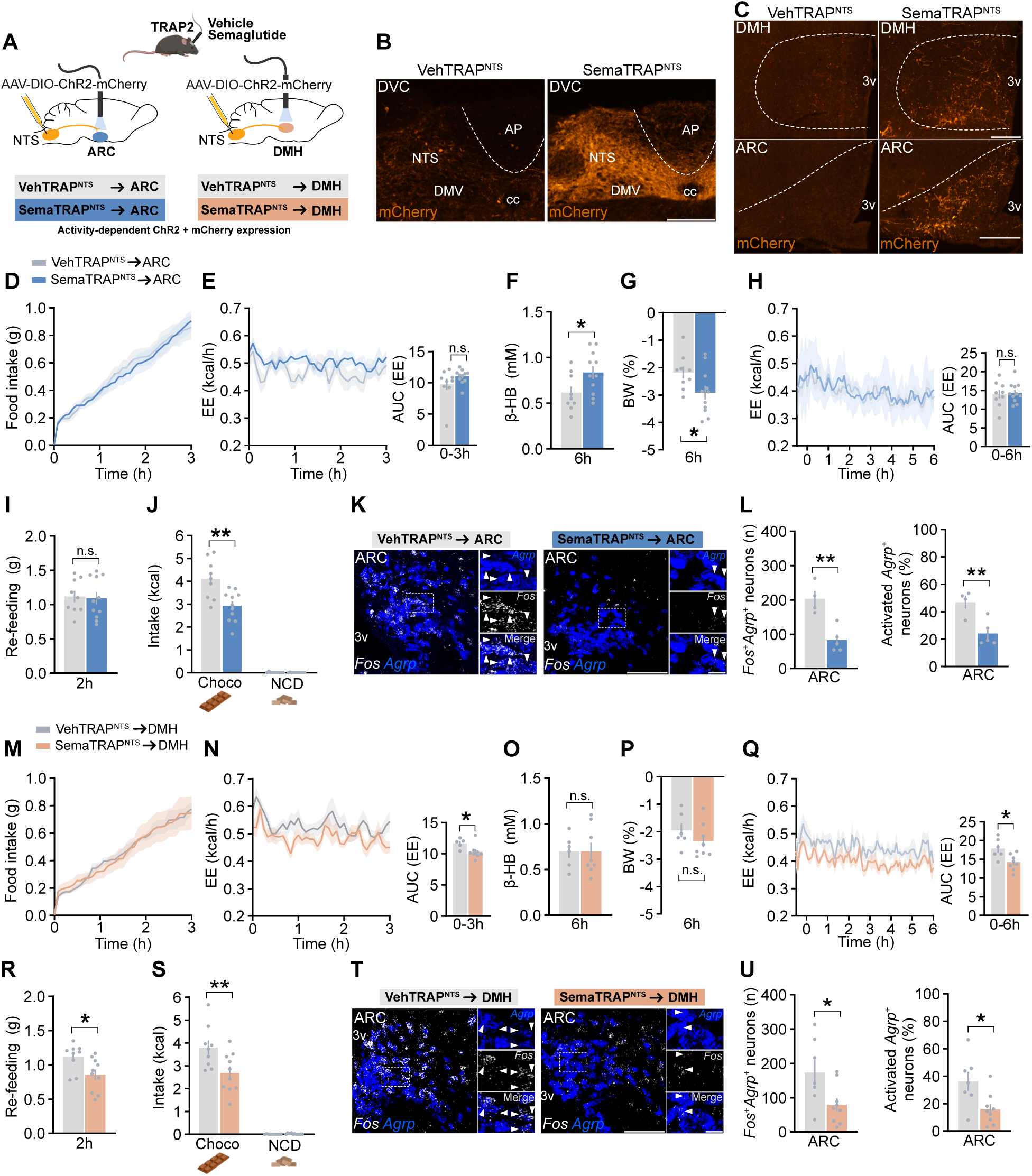
Semaglutide-responsive NTS→hypothalamus projections are functionally specialized for motivated feeding and metabolic control. (A) Approach to target semaglutide-responsive NTS-derived projections to the ARC and DMH using TRAP2 technology in combination with optogenetics. (B, C) Expression of mCherry in the NTS (B) and in NTS-derived projections in the ARC and DMH (C) of vehTRAP^NTS^ mice (reflecting background TRAP) and semaTRAP^NTS^ mice (reflecting semaglutide-responsive neurons and their projections). (D, E) Effect of photostimulation on normal dark phase feeding and energy expenditure in semaTRAP^NTS^**→**ARC mice, compared to that in vehTRAP^NTS^**→**ARC mice. *n=9-11*. (F-H) Effect of photostimulation on ketogenesis (F), body weight (G) and energy expenditure (H) in semaTRAP^NTS^**→**ARC compared to that in vehTRAP^NTS^**→**ARC mice, during the light phase in the absence of feeding. *n=9-11*. (I, J) Effect of photostimulation on fasting-induced refeeding (I) and on binge-eating of chocolate and concomitant intake of normal chow during a 2 h restricted access paradigm (J) in semaTRAP^NTS^**→**ARC mice, compared to that in vehTRAP^NTS^**→**ARC mice. *n=9-11*. (K) Fluorescent *in situ* hybridization showing *Agrp* (blue) and *Fos* (white) mRNA expression in the ARC of vehTRAP^NTS^**→**ARC and semaTRAP^NTS^**→**ARC mice fasted for 4 h into the dark phase with concomitant photostimulation. (L) Quantification of co-expression in (K). *n=4-5*. (M, N) Effect of photostimulation on normal dark-phase feeding (M) and energy expenditure (N) in semaTRAP^NTS^**→**DMH mice, compared to that in vehTRAP^NTS^**→**DMH mice. *n=5-8*. (O-Q) Effect of photostimulation on ketogenesis (O), body weight (P) and energy expenditure (Q) in semaTRAP^NTS^**→**DMH compared to that in vehTRAP^NTS^**→**DMH mice, during the light phase in the absence of feeding. *n=6-8*. (R, S) Effect of photostimulation on fasting-induced refeeding (R) and on binge-eating of chocolate and concomitant intake of normal chow during a 2 h restricted access paradigm (S) in semaTRAP^NTS^**→**DMH mice, compared to that in vehTRAP^NTS^**→**DMH mice. *n=9-10*. (T) Fluorescent *in situ* hybridization showing *Agrp* (blue) and *Fos* (white) mRNA expression in the ARC of vehTRAP^NTS^**→**DMH and semaTRAP^NTS^**→**DMH mice fasted for 4 h into the dark phase with concomitant photostimulation. (U) Quantification of co-expression in (T). *n=5-7*. n.s.=not significant, \**p*<0.05, \*\**p*<0.01. Data were analyzed using unpaired t-test (E-J, L, N-S, U) or two-way ANOVA with Šidák’s post-hoc test (D, M). Error bars represent ± SEM. Scale bars = 200 µm (B, C), 100 µm (K, T) and 20 µm in magnified boxed regions (K, T). Arrowheads point at double-labeled neurons. AP, area postrema; ARC, arcuate nucleus; cc, central canal; DMH, dorsomedial hypothalamus; DMV, dorsal motor nucleus of the vagus; DVC, dorsal vagal complex; NTS, nucleus of the solitary tract; 3v, third ventricle. Relates to Figure S4.

Under normal dark-phase conditions, activation of the semaTRAP^NTS^→ARC projection did not alter food intake or EE (Figures 6D and 6E) or RER (Figure S4A) compared to controls. Because *Adcyap1*^AP/NTS^ neurons are required for semaglutide-induced ketogenesis during the inactive phase^11^, we next asked whether photostimulation of the semaglutide-responsive NTS→ARC projection is sufficient to reproduce this effect. Indeed, specific stimulation of the semaTRAP^NTS^→ARC projection during the inactive phase, in the absence of feeding, promoted ketogenesis and weight loss compared to vehTRAP^NTS^ controls (Figures 6F and 6G), while effects on EE and RER did not reach statistical significance under these conditions (Figures 6H and S4B).

Semaglutide suppresses not only homeostatic feeding but also intake of palatable food^11,29,30^. Notably, stimulation of the semaTRAP^NTS^→ARC projection failed to alter fasting-induced refeeding, but suppressed binge-like chocolate consumption (Figures 6I and 6J). Moreover, fasting-induced activation of AgRP neurons was significantly suppressed by stimulation of semaTRAP^NTS^→ARC projections (Figures 6K and 6L). Finally, there was a strong trend towards suppression of overall *Fos* mRNA expression in the ARC of semaTRAP^NTS^→ARC mice compared to vehTRAP^NTS^ control mice (Figure S4C).

Specific stimulation of the semaTRAP^NTS^→DMH projection under normal dark-phase conditions did not alter food intake or RER (Figures 6M and S4D) but significantly lowered EE (Figure 6N), mirroring the effects observed following stimulation of projections from the broader *Adcyap1*^NTS^ population. During the inactive phase, stimulation of the semaTRAP^NTS^→DMH projection did not affect ketogenesis, body weight or RER (Figures 6O, 6P and S4E), but did significantly lower EE (Figure 6Q), consistent with the effect observed under free-feeding conditions (Figure 6N).

Stimulation of the semaTRAP^NTS^→DMH projection reliably suppressed both fasting-induced refeeding and chocolate binge-eating (Figures 6R and 6S), reproducing the context-dependent suppression of motivated feeding observed following stimulation of projections from the broader *Adcyap1*^NTS^ population. Fasting-induced activation of AgRP neurons was likewise significantly suppressed by stimulation of semaTRAP^NTS^→DMH projections (Figures 6T and 6U). Moreover, we confirmed that photostimulation under these conditions reliably induced c-Fos expression in the DMH of semaTRAP^NTS^→DMH mice compared to vehTRAP^NTS^→DMH controls (Figure S4F).

Together, these findings demonstrate that stimulation of semaglutide-responsive NTS→ARC and NTS→DMH projections reproduce key projection-specific behavioral and metabolic phenotypes observed following activation of projections arising from the *Adcyap1*^NTS^ population. These results therefore identify distinct semaglutide-responsive brainstem-to-hypothalamus pathways that engage the ARC and DMH to suppress AgRP neuron activity and differentially regulate motivated feeding and metabolic state.

## DISCUSSION

Here, we identify previously unrecognized *Adcyap1*^NTS^ brainstem-to-hypothalamus pathways that differentially regulate motivated feeding and metabolism. Rather than representing a uniform satiety output, *Adcyap1*^NTS^ projections to the ARC and DMH exhibit striking functional specialization. Both pathways selectively suppress feeding during states of elevated motivational drive while sparing physiological dark-phase feeding, yet they exert distinct metabolic effects: whereas the *Adcyap1*^NTS^→ARC pathway promotes ketogenesis and weight loss independently of food intake, the *Adcyap1*^NTS^→DMH pathway selectively reduces energy expenditure. Both pathways converge on suppression of fasting-induced activation of AgRP neurons. Furthermore, selective activation of semaglutide-responsive NTS projections to the same hypothalamic targets recapitulated key projection-specific functions of the corresponding *Adcyap1*^NTS^ pathways, demonstrating that semaglutide recruits NTS→ARC and NTS→DMH pathways capable of exerting distinct aspects of its effects. These findings thus establish a framework through which semaglutide suppresses highly motivated feeding and regulates metabolic state.

### Brainstem-mediated semaglutide control of AgRP neurons

Previous work has identified ARC and DMH neuronal populations capable of mediating part of GLP-1RA-induced anorexia and weight loss^6–8,31^. These populations are largely GABAergic and inhibit AgRP neurons^6,7^, consistent with earlier *ex vivo* studies showing that GLP-1RAs indirectly suppress AgRP neurons, which themselves express little to no GLP-1R^32–34^. The contribution of local hypothalamic GLP-1R circuitry to GLP-1RA-induced weight loss, however, remains incompletely resolved. Silencing the TRH-expressing ARC neurons, which express GLP-1R, attenuates liraglutide-induced weight loss^6^, whereas ablation of ARC GLP-1R-expressing neurons leaves semaglutide-induced weight loss intact^10^. Moreover, whether the indirect suppression of AgRP neurons by GLP-1RAs is mediated solely by local action on circuits within the hypothalamus or also through upstream GLP-1RA-responsive inputs has remained unknown. The DVC is a potential source of such input and a major site of semaglutide action^10–13,35^, with GLP-1R^+^ neurons in the AP implicated as upstream drivers of both non-aversive and aversive GLP-1RA effects^10,12^. Our recent work placed *Adcyap1*^NTS^ neurons downstream of AP GLP-1R signaling^11^, but how this DVC-derived signal reaches hypothalamic circuits controlling feeding and metabolism remained unresolved. Our findings now identify a previously unrecognized DVC-to-hypothalamus route through which semaglutide can suppress fasting-induced AgRP neuron activation, as activation of semaglutide-responsive NTS→ARC and NTS→DMH projections each reproduced this effect. This DVC-derived control of AgRP neurons aligns with recent *in vivo* fiber photometry showing that exendin-4 rapidly suppresses AgRP neuron activity in fasted mice and attenuates the subsequent response to food presentation, with the magnitude of this modulation predicting the reduction in food intake^9^. Consistent with a functional contribution of AgRP neuron suppression, chemogenetic activation of AgRP neurons under our DIO subchronic treatment paradigm reversed semaglutide-induced reductions in food intake and body weight. Recent work further suggests that the contribution of AgRP neurons to semaglutide action is context-dependent, with intact AgRP neuron function required for sustained weight loss under specific experimental conditions^36^. Together, these findings indicate that suppression of AgRP responses can contribute to the feeding effects of semaglutide, while the role of AgRP neurons may differ during sustained treatment.

At the level of the DVC relay, our data show that semaglutide recruits a transcriptionally heterogeneous *Adcyap1*^NTS^ population. While endotoxin-responsive *Adcyap1*^NTS^ neurons are glutamatergic^21^, we find that approximately one third of *Adcyap1*^NTS^ neurons, and about half of the semaglutide-activated subset, express *Slc32a1*, indicating an inhibitory phenotype. While we did not determine whether the *Adcyap1*^+^ projections to the ARC or DMH were excitatory or inhibitory, the presence of both excitatory and inhibitory phenotypes within the *Adcyap1*^NTS^ population is compatible with our optogenetic data in mice fasted into the dark phase, showing that photostimulation gave rise to divergent effects in the two target regions, with NTS→DMH stimulation increasing c-Fos expression in the DMH, whereas NTS→ARC stimulation was associated with a trend toward reduced c-Fos expression in the ARC.

### Semaglutide-responsive NTS pathways spare physiological dark-phase feeding

A recently described GABAergic NTS→ARC pathway, not examined in the context of GLP-1RA action, suppresses dark-phase chow intake non-aversively^24^. In contrast, the *Adcyap1*^NTS^-derived pathways examined here spare normal dark-phase feeding despite suppressing fasting-induced refeeding, suggesting that hypothalamus-projecting NTS populations can regulate distinct components of feeding behavior. The absence of an effect on normal dark-phase intake may reflect state-dependent differences in both the regulation and functional contribution of AgRP neurons. AgRP neuron activity exhibits a circadian rhythm that peaks around feeding onset and is regulated by SCN→DMH→ARC circuitry^16,37^, whereas ablation of AgRP/NPY neurons in adult mice leaves *ad libitum* feeding largely intact while blunting fasting-induced refeeding^38^. Thus, *Adcyap1*^NTS^ input may preferentially counter fasting-related AgRP neuronal drive without being sufficient to disrupt the circadian and parallel circuits supporting normal dark-phase feeding. This interpretation is also consistent with our previous finding that ablation of *Adcyap1*^AP/NTS^ neurons does not affect baseline feeding or body weight in lean mice^11^.

### Non-aversive brainstem-to-hypothalamus pathways in semaglutide control of motivated feeding

The specific ability of *Adcyap1*^NTS^→ARC and *Adcyap1*^NTS^→DMH projections to suppress fasting-induced refeeding and binge-like chocolate intake identifies a brainstem-derived mechanism that restrains feeding under conditions of elevated motivational drive. Although the NTS has classically been viewed as a regulator of short-term homeostatic feeding, our findings contribute to an evolving view of the DVC as a more diverse regulator of energy balance, and that such effects on energy balance are not necessarily due to aversive signaling^39,40^. Importantly, stimulation of these pathways did not induce CTA, arguing against an aversive mechanism underlying their feeding effects. Together with our previous finding that semaglutide-induced CTA is preserved following ablation of *Adcyap1*^AP/NTS^ neurons^11^, these data suggest that *Adcyap1*^NTS^-derived hypothalamic pathways preferentially contribute to a non-aversive component of semaglutide action. Selective activation of semaglutide-responsive NTS→ARC and NTS→DMH projections further linked this control of motivated feeding to drug-responsive pathways, with both projections suppressing chocolate binge-eating. Together with recent evidence implicating CeA GLP-1R neurons in GLP-1RA control of reward-driven feeding^29^, our findings indicate that GLP-1RAs can recruit anatomically distinct central circuits to restrain different aspects of palatable-food intake.

Fasting-induced refeeding was reduced following activation of the semaglutide-responsive NTS→DMH, but not NTS→ARC, projection. Because other NTS→ARC responses were preserved, this difference is unlikely to reflect a general failure to recruit the pathway and may instead reflect a function of a broader *Adcyap1*^NTS^→ARC pathway relative to a more restricted semaglutide-responsive projection.

### Metabolic specialization of NTS-to-hypothalamus pathways

We previously identified a DVC-dependent component of semaglutide-induced ketogenesis and weight loss that occurs independently of caloric intake^11^. In the present study, activation of both *Adcyap1*^NTS^→ARC and semaglutide-responsive NTS→ARC projections similarly induced ketogenesis and weight loss, localizing a component of this DVC-dependent metabolic response to a defined brainstem-to-hypothalamus pathway. The convergence of these independently targeted NTS→ARC projections on the same metabolic phenotype further supports the NTS→ARC route as a functionally relevant component of the semaglutide response. AgRP neurons have an established role in regulating peripheral substrate utilization^28^ and were recently shown to regulate fasting-associated hepatic metabolism, with direct optogenetic AgRP neuron activation promoting β-oxidation and ketone-body production and AgRP neuron inhibition during fasting reducing hepatic ketogenesis^41^. These findings argue against a simple model in which inhibition of AgRP neurons mediates the ketogenic effect of NTS→ARC stimulation. Indeed, activation of both the NTS→ARC and the NTS→DMH pathway reduces fasting-induced AgRP neuron activation, whereas only the NTS→ARC projection promotes ketogenesis. However, our *Fos* mRNA measurements were obtained after a short fasting period at dark-phase onset, whereas ketogenesis was assessed during the inactive phase in the absence of feeding. Thus, these experiments cannot establish a direct relationship between AgRP response and the ketogenic phenotype. Moreover, whereas Chen et al. manipulated AgRP neurons directly, our experiments targeted an upstream NTS afferent to the ARC. Our data therefore establish NTS→ARC activation as sufficient to promote ketogenesis and weight loss, while leaving the downstream ARC mechanism underlying this metabolic response unresolved. Together with our previous finding that ablation of *Adcyap1*^AP/NTS^ neurons suppresses semaglutide-induced ketogenesis and preferentially attenuates semaglutide-induced fat loss^11^, these results reinforce a role for *Adcyap1*^AP/NTS^ circuitry in semaglutide-induced metabolic responses that are dissociable from reduced caloric intake.

In contrast, activation of *Adcyap1*^NTS^→DMH and semaglutide-responsive NTS→DMH projections reduced energy expenditure while sparing active-phase chow intake, dissociating this metabolic effect from an acute change in feeding. The DMH contains functionally heterogeneous neurons capable of regulating energy expenditure in opposing directions, with previous studies showing that distinct DMH populations and afferent pathways can either increase thermogenesis and energy expenditure or promote hypometabolic states^42–44^. Our findings therefore identify a semaglutide-responsive NTS→DMH input capable of driving an energy-conserving output, distinct from the ketogenic and weight-lowering effects of NTS→ARC pathway activation.

## Conclusion

Collectively, our findings identify previously unrecognized *Adcyap1*^NTS^→ARC and *Adcyap1*^NTS^→DMH pathways that differentially regulate highly motivated feeding and metabolic state, while semaglutide-responsive NTS projections to the same hypothalamic targets reproduce key pathway-specific functions. By functionally interrogating semaglutide-responsive NTS projections to the ARC and DMH, our study therefore identifies how a major relay for semaglutide effects, the DVC, engages two hypothalamic regions central to energy balance and distinguishes separable behavioral and metabolic components of the drug’s action. Further resolution of GLP-1RA-responsive circuitry may ultimately identify targets that can be selectively engaged, providing a basis for more precisely designed anti-obesity therapies and a more complete understanding of the neural control of energy balance.

## Limitations of the study

Our study has several limitations that should be considered when interpreting these circuit effects. The experiments were not designed to assess sex-dependent differences in circuit function. In addition, optogenetic stimulation was performed unilaterally, which may have limited sensitivity to detect modest effects under conditions in which baseline values were already low. Our projection-specific manipulations primarily establish sufficiency and therefore define the functional capacity of semaglutide-responsive NTS→ARC and NTS→DMH pathways rather than their relative contribution to the complete drug response. Moreover, activation of AgRP neurons was assessed early in the dark phase, whereas several metabolic phenotypes, including ketogenesis, were examined during the inactive phase; thus, the observed effects on AgRP neurons cannot necessarily be extrapolated across physiological states or assumed to mediate the metabolic responses.

## ACKNOWLEDGMENTS

The Centre for Cellular Imaging at the University of Gothenburg is acknowledged for their support. Some images were created with icons or templates from BioRender.com. L.E.R. received funding from the Novo Nordisk Foundation (NNF21OC0067317, Excellence Emerging Investigator Grant - Endocrinology and Metabolism 2021), the Swedish Research Council Starting Grant 2020 (Medicine and Health; 2020-02473), the Swedish Research Council Consolidator Grant 2024 (Medicine and Health; 2024-03045), the Swedish Brain Foundation (FO2024-0376) and the Sahlgrenska Academy (2022/334). J.R. received funding from the Swedish Research Council (2017-01611 and 2025-03430).

## AUTHOR CONTRIBUTIONS

Conceptualization, project administration, supervision and funding acquisition: L.E.R.

Investigation: S.B.S, J.T.D. and L.E.R

Formal analysis and visualisation: S.B.S, J.T.D. and L.E.R.

Writing: S.B.S and L.E.R, with input from J.T.D and J.R.

Resources: J.R. and L.E.R.

## METHODS

### Animals

Mice were housed on a 12:12h light:dark cycle (lights on at 7 am, lights off at 7 pm) at an ambient temperature of approximately 21°C with ad libitum access to food and water unless stated otherwise. During breeding, mice were fed a Teklad global complete feed (#2018C, Envigo, Madison WI, USA) while from weaning and onwards, a Teklad global diet (#2916) was provided. For diet-induced obesity experiments, mice were given a high fat diet (60% kcal from fat; #E15742-340, Ssniff) from the age of 7-10 weeks and onwards. All experiments were carried out in adult male mice. Prior to experiments, mice were single housed for at least 7 days to acclimate to the experimental condition. All animal procedures were carried out at the Experimental Biomedicine facility at the University of Gothenburg, Sweden, and followed national and European guidelines. All procedures had been approved by the Animal Care and Use Committee in Gothenburg, Sweden.

### Mouse lines

All mice were purchased from Jax and included: *Agrp^tm1(cre)Lowl^*/J (AgRP-Ires-Cre; #012899) ^45^, Fos^tm2.1(icre/ERT2)Luo^/J (TRAP2 mice; #030323)^46^, B6N;129-Tg(CAG-CHRM3*,-mCitrine)1Ute/J (R26-hM3Dq/mCitrine; #026220)^47^, *Adcyap1^tm1.1(cre)Hze^*/ZakJ (*Adcyap1*-2A-Cre: #035239)^48^ and C57BL/6J mice (Charles River, Germany). Heterozygous TRAP2 and *Adcyap1*-2A-Cre mice were generated by crossing tg/tg mice with C57BL/6J mice. AgRP-Ires-Cre tg/wt mice were crossed with double-floxed R26-hM3Dq mice, generating Cre-positive and Cre-negative experimental mice, heterozygous for the hM3Dq allele.

### Viral vectors and stereotactic surgery

All viral vectors were purchased from Addgene and included; for chemogenetic experiments, AAV5 or AAV8-hSyn-DIO-hM3D(Gq)-mCherry and AAV5 or AAV8-hSyn-DIO-mCherry^19^ (control); for neuronal Cre-dependent ablation studies, AAV5-EF1a-DIO-taCasp3-TEVp^20^ and AAV5-hSyn-DIO-mCherry^19^ (control); for optogenetic experiments, AAV5-EF1a-DIO-hChR2(H134R)-mCherry-WPRE-HGHpA. pAAV-hSyn-DIO-hM3D(Gq)-mCherry and pAAV-hSyn-DIO-mCherry were a gift from Bryan Roth (Addgene viral preps #44361-AAV8 and #44361-AAV5, #50459-AAV8 and #50459-AAV5), pAAV-EF1a-flex-taCasp3-TEVp was a gift from Nirao Shah & Jim Wells (Addgene viral prep #45580-AAV5) and pAAV-EF1a-double floxed-hChR2(H134R)-mCherry-WPRE-HGHpA was a gift from Karl Deisseroth (Addgene viral prep # 20297-AAV5).

For stereotactic surgeries, mice were anesthetized with isoflurane and placed in a stereotaxic frame. For targeting the DVC, the coordinates relative to bregma were: anteroposterior (AP) −7.5 mm, mediolateral (ML) ±0.2 mm and dorsoventral (DV) −4.4 and −4.3 mm. The skull was exposed, two drill holes were made, and bilateral injections were performed (75 nl per depth for Gq-viruses and ChR2; 25 nl per depth for Casp3-virus; 5 nl/s) using a Nanoject III automated injector (#3-000-207, Drummond Scientific). The skin was sutured, and the mice were placed in a new cage after the procedure and given at least 3-4 weeks of recovery before being used in any experiment. Virus expression was verified histologically in all animals after sacrifice. In mice destined for optogenetics, a flat tip fiber-optic cannula [5 mm (DMH) or 6 mm (ARC), 200 µm in diameter, NA 0.37, Doric Lenses Inc.] was implanted above above the ARC (coordinates relative to Bregma: AP −1.6, ML −0.2, DV −5.4) or the DMH (coordinates relative to Bregma: AP −1.6, ML −0.3, DV −4.7) immediately after the viral vector delivery, and secured to the skull via dental cement (SuperBond C&B).

### Drugs and administration

Semaglutide (Ozempic^®^, Novo Nordisk), was diluted in sterile PBS prior to experiments and injected s.c. at a dose of 15 or 60 μg/kg body weight based on previous studies^18,29^. For subchronic experiments with daily injections, the dose was titrated up to 60 μg/kg: 15 μg/kg on day 1, 30 μg/kg on day 2, and then 60 μg/kg for the remainder of the experiment. CNO (#C0832, Sigma Aldrich) was dissolved in dimethyl sulfoxide (DMSO) and administered i.p at a dose of 0.1 mg/kg in chemogenetic experiments involving TRAP2 mice or *Adcyap1*-2A-Cre mice and 1 mg/kg for experiments with AgRP-IRES-Cre mice, diluted in sterile 0.9% w/v saline. Ghrelin (#1465; Tocris, Bristol, UK) was dissolved in sterile saline and injected i.p at a dose of 1 mg/kg.

### TRAP procedure

TRAP2 mice, stereotactically injected with AAV8-hSyn-DIO-hM3D(Gq)-mCherry, AAV8-hSyn-DIO-mCherry or AAV5-EF1a-DIO-hChR2(H134R)-mCherry-WPRE-HGHpA in the DVC, were given 3-4 weeks to recover from surgery and to allow for viral expression. On the TRAP day, the mice had their food removed and received a s.c. injection of vehicle (PBS) or semaglutide (60 μg/kg for subsequent chemogenetics, 120 μg/kg for subsequent optogenetics), followed by i.p. injection of 4TM (20 mg/kg) 4 hours later. 4TM was prepared in an aqueous solution as previously described^49^. The mice were then given 2-3 weeks for expression of the AAV-encoded hM3Dq and/or mCherry before any experiments were conducted.

### Manual food intake recordings after drug administration

To investigate the impact of semaglutide on ghrelin-induced feeding, food was withdrawn at the beginning of the light phase and vehicle or semaglutide (60 μg/kg body weight, s.c.) was administered. 4 hours later, vehicle or ghrelin (1 mg/kg body weight, i.p.) was injected, and normal chow was returned. Food intake was then manually measured at 2, 4 and 6 hours. For fasting–refeeding experiments, mice were given a clean cage and food was removed just before the onset of the dark phase (19:00). After an overnight fast, CNO (0.1 mg/kg, i.p.), vehicle, or semaglutide (60 μg/kg, s.c.) was administered, CNO immediately prior to light onset (when food was returned) and vehicle or semaglutide 1 hour before light onset. Food intake and body weight were manually measured 2 hours later. For subchronic experiments in DIO mice, mice were on a high fat diet for approximately 12 weeks before receiving daily injections with vehicle or semaglutide 3 hours before the dark phase. Food intake and body weight were measured daily. When combined with CNO treatment, CNO was injected i.p. just prior to the dark phase.

### *In vivo* optogenetics

Prior to any experiments, mice were habituated to being connected to a fiberoptic patch cord (core diameter 200 µm, NA 0.22; Doric Lenses Inc.) secured to a rotary joint above the cage (Doric Lenses). The light source was a 450 nm laser and the photostimulation protocol was composed of repeated sequences of 10 ms light pulses at a frequency of 30 hz for 1 second, followed by a 4-second break^24^ (programmed in Doric Neuroscience Studio). The output power was 20 mW at the tip of the fiber. All optogenetic experiments in *Adcyap1*-2A-Cre mice except for the conditioned taste aversion protocol were performed in a cross-over design, where half of the mice received photostimulation (laser ON) and half of the mice were connected to patch cords but received no photostimulation (laser OFF). The experiment was then reversed 4-7 days later. For optogenetic experiments in TRAP2 mice, where mice TRAPed with vehicle served as controls, all mice received photostimulation. Optic fiber placement and AAV expression were verified in all mice after sacrifice.

#### Automated food intake and indirect calorimetry

For optogenetic experiments with concurrent indirect calorimetry, mice were housed in Promethion Core metabolic cages (Sable Systems). The Promethion Core^TM^ CGF (Combined Gas Analysers and Flow Regulation) records gas flow and composition and automatically calculates RER as the ratio between VCO_2_ and VO_2_, and EE according to the Weir equation. These cages were equipped with FED3.1 (Feeding Experimentation Device version 3.1), an open-source home-cage compatible feeding device, set to free-feeding mode. The FED was placed outside of the cage, but made accessible through a small customized opening in the cage. The FEDs dispense a 20-mg chow pellet (#1811142, Operant Chamber Resources, AIN-76A LabTab Rodent Tablet, OCB Systems, UK) into the feeding well of the device, monitored by a beam. The beam breaks when mice remove the pellet, and the FED logs the timestamp and dispenses a new pellet, allowing for continuous food intake recordings. Mice were habituated to the cages and the FEDs for at least 3-4 days prior to any experiments. For dark phase feeding experiments in the Promethion Core system. On the day of the experiment, mice were connected to a patch cord 3 hours before dark onset with the FEDs turned off. Photostimulation was initiated 15 minutes before dark onset, and the FEDs were switched on just prior to lights-off. Photostimulation was allowed to continue for 3 h. The food intake data was analyzed using an open-source software FED3VIZ^27^.

For calorimetry recordings in the absence of feeding, mice received a clean cage on the day before the experiment. On the following day, one hour after light phase onset, mice were weighed and connected to patch cords. After 30 minutes, photostimulation was initiated and allowed to continue for 6 hours. Mice were again weighed and a small blood sample was taken from the tail tip, and blood ketone levels were measured using test strips and a ketone meter (Keto-Mojo, Amsterdam-Duivendrecht, Netherlands).

#### Manual food/chocolate intake recordings

For fasting-refeeding experiments, mice were fasted overnight. At light onset, mice were weighed and connected to patch cords and photostimulation was initiated. After 15 min of stimulation, normal chow was again made available and mice were allowed to eat for 2 h, after which both food intake and body weight were recorded.

To assess hedonic feeding, freely fed mice were given access to chocolate (Marabou milk chocolate, ∼5 kcal/g; Mondelez Sverige AB) alongside normal chow for 2 hours during the light phase (11.00 – 13.00) for 5 consecutive days to establish entrained and stable chocolate intake. On day 6, mice were connected to patch cords 1 hour prior to chocolate access, and photostimulation was applied during the 2-hour access period. Intake of normal chow and chocolate was measured manually. For optogenetic experiments in *Adcyap1*-2A-Cre mice, a laser ON/OFF paradigm was employed. Half of the mice received photostimulation during the chocolate session on day 6, while the remaining mice served as unstimulated controls. On day 7, all mice underwent the chocolate session without photostimulation. On day 8, photostimulation was applied to the mice that had not received stimulation on day 6.

#### Conditioned taste aversion (CTA)

The CTA protocol was adapted from Chen et al^50^. The mice were deprived of water just before dark phase onset on day 0. On day 1-3, the mice had access to two non-leaking water bottles (Drinko Measurer, leak-free sipper tube, Amuza Inc, USA) during two daily time-windows: 11.00– 11.30 (AM) and 16.30–17.30 (PM). This scheduled drinking ensured that the mice were thirsty and ready to drink as soon as bottles were available. On day 4 (conditioning day), mice were connected to patch cords one hour before getting access to a 5% sucrose solution in both bottles. After 30 min of access, half of the mice received 1 h of photostimulation while the other half did not receive any stimulation (laser ON vs. laser OFF for optogenetics in the Adcyap1-2A-Cre line; laser ON for all mice when stimulation was performed in TRAP2 mice). After stimulation, mice were disconnected from the patch cords and were again given water in the afternoon. On day 5, mice followed the same drinking schedule as on day 1-3. On day 6 (test day), mice were given a choice between one bottle with water and one with 5% sucrose solution for 30 minutes. Bottle positions were randomly distributed to control for any side preference. Sucrose preference was calculated as the amount of sucrose solution consumed divided by the total fluid intake, measured in grams.

### Tissue preparation

Mice were anesthetized with a mixture of medetomidine and ketamine i.p. and were, after confirmation of sedation by a reflex test, perfused transcardially; first using room-tempered 0.9% saline, followed by ice-cold paraformaldehyde (4% PFA, pH 7.4, dissolved in 1X phosphate-buffered saline). Brains were carefully extracted and post-fixed in 4% PFA solution for approximately 20 h at 4°C. In the case of optogenetics, whole heads were post-fixed in 4% PFA solution for approximately 16h at 4°C in order to preserve fiber placement. Brains were then carefully removed from the skulls and post-fixed for an additional 4 hours in 4% PFA. After post-fixation, the brains were cryoprotected by submersion in 25% sucrose 1X PBS solution for at least 24h. The brains were then cut at 20 µm on a freezing sledge microtome (Bright Instruments, UK). Sections were stored free-floating at −20°C in an antifreeze solution (25% ethylene glycol; 25% glycerol; 0.05M phosphate buffer) until further use.

### Fluorescent immunohistochemistry

All steps were carried out at room temperature. Sections were incubated in a blocking solution (1% bovine serum albumin and 0.3% Triton X-100 in 1X PBS) for 45 minutes, followed by overnight incubation in primary antibodies (guinea pig anti-c-Fos, 1:5000; Synaptic Systems) diluted in blocking solution. For co-detection of virus expression and projections (mCherry), an anti-tdTomato (1:500; Sicgen) primary antibody was added. Sections were rinsed in 1X PBS for 30 minutes and subsequently incubated in a secondary antibody solution for 1 h at room temperature: HRP-conjugated donkey anti-guinea pig (1:2000, Invitrogen) for c-Fos detection; donkey anti-sheep 555 (1:500, Invitrogen) for mCherry detection, diluted in blocking solution. Sections were then rinsed in 1X PBS for 30 minutes. For c-Fos detection, the sections were incubated in tyramide solution (Alexa Fluor^TM^ 488 Tyramide SuperBoost^TM^ kit 1:200; Invitrogen) for 10 minutes and then rinsed in 1X PBS for 30 minutes. For visualizing brain-areas and orientation, a red or blue fluorescent Nissl stain was sometimes employed: 20 minutes incubation in NeuroTrace 435/455 or 530/615 (1:100; Invitrogen), followed by a 10-minute rinse in 0.5% Tween-20 1X PBS solution. Lastly, the sections were rinsed in 1X PBS for at least 1h before they were mounted on SuperFrost Plus slides and covered with ProLong Gold antifade mounting medium (Invitrogen).

### Imaging and quantification of immunohistochemistry

Images were captured with an epi-fluorescent wide-field microscope (AxioObserver Z1, Zeiss), using a 20x objective and a 14 bit AxioCam 506 mono camera (Zeiss). The ARC and the DVC were imaged at three to four and two different rostro-caudal levels, respectively using tile scans. The images were stitched and shading correction was applied using the Zen Blue software (v.2.6; Zeiss) before being imported into ImageJ/FIJI (NIH, Bethesda), where brightness and contrast were adjusted equally over each channel. Regions of interest (ROIs) were then drawn around the area to be analysed, whereafter c-Fos positive neurons were counted manually using the Cell Counter plug-in.

### Fluorescent *in situ* hybridization

Fluorescent *in situ* hybridization using RNAscope (ACD; Advanced Cell Diagnostics Inc., Hayward, CA, USA) was performed to simultaneously detect *Agrp* and *Fos* mRNA in the ARC, or *Adcyap1*, *Slc32a1* and *Fos* mRNA in the DVC. The *Agrp* probe (#400711;) contained 16 oligonucleotide pairs and targeted region 11-764 (Acc. no. NM_001271806.1) of the *Agrp* transcript, the Adcyap1 probe (# 405911) contained 20 oligo pairs and targeted region 676-1859 of the *Adcyap1* mRNA (Acc. no. NM_009625.2), the *Slc32a1* probe (#319191) contained 20 oligo pairs and targeted region 894-2037 of the *Slc32a1* transcript (Acc. no. NM_009508.2), while the *Fos* probe (#316921) contained 20 oligonucleotide pairs and targeted region 407-1427 (Acc. No. NM_010234.2) of the *Fos* transcript. All reagents were obtained from ACD. The pre-treatment protocol followed the manufacturer’s instructions, including target retrieval for 8 min at 98°C and a 15-30 min incubation with Protease Plus. For detection, the RNAscope Multiplex Fluorescent v2 kit was used according to manufacturer’s instructions. Opal 570 was used for *Fos* mRNA detection (1:2000; Akoya Biosciences) and Opal 520 for detection of *Agrp or Adcyap1* mRNA (1:500 and 1:2000, respectively). For the detection of *Slc32a1* or *Adcyap1* mRNA in ablated mice, Opal 650 was used (1:2000).

### Imaging and quantification of *in situ* hybridization

For quantification of RNAscope data, images were captured using a laser scanning confocal microscope (LSM 700 inverted; Zeiss) equipped with a Plan-Apochromat 40x/1.3 Oil objective. Images were captured using tile scans with Z-stacks. For the ARC, three to four unilateral or bilateral (ablation studies) images per animal were captured along the rostro-caudal extent of the ARC, for the DVC, two images were captured per animal. Laser intensities were kept constant throughout the imaging process for the respective cohorts. The raw images were stitched using the Zen Black software (Zeiss) and imported into ImageJ/FIJI for conversion to maximum intensity projections. The channels were merged, and the images were imported into an open-source software, QuPath^51^ (v.0.5.1), for quantification purposes. The software relies on DAPI for cell recognition and quantifies the number of dots/cell. The number of dots, including estimated dots from clusters and single dots, were used to classify neurons as positive/negative for the specific mRNA transcript analyzed. Depending on the probe and fluorophore, a minimum threshold of 5-8 estimated dots, corresponding to mRNA molecules, was used.

### Statistical analysis

Data are reported as mean values and error bars represent standard error of the mean (±SEM). For comparison of one variable between two groups, an unpaired, two-sided Student’s t-test was used, except for the optogenetic experiments in which each mouse was its own control, in which a paired t-test was employed. For c-Fos counts in multiple structures, multiple unpaired t-tests were performed, with Holm-Šídák’s *post hoc* test to correct for multiple comparisons. Comparisons between several independent datasets or groups were performed using one-way ANOVA followed by Tukey’s or Šídák’s *post hoc* test. For datasets with multiple factors/variables, a two-way ANOVA followed by Tukey’s or Šídák’s post hoc test was used. All statistical analyses were performed using GraphPad Prism v.10.2.0 (GraphPad; La Jolla, CA). Significance level was set to a p value lower than 0.05 (p<0.05). Differences are marked with */# or ¤, indicating ^*/#^p<0.05, ^**/##/¤¤^p<0.01, ^***/###^p<0.001 or ^****/####/¤¤¤¤^p<0.0001 respectively, as specified in the figure legends.

## Declaration of generative AI and AI-assisted technologies in the manuscript preparation process

During the preparation of this work, the authors used ChatGPT for language improvement. The authors reviewed and edited the output as needed and take full responsibility for the content of the published article.

---

**Figure S1:**
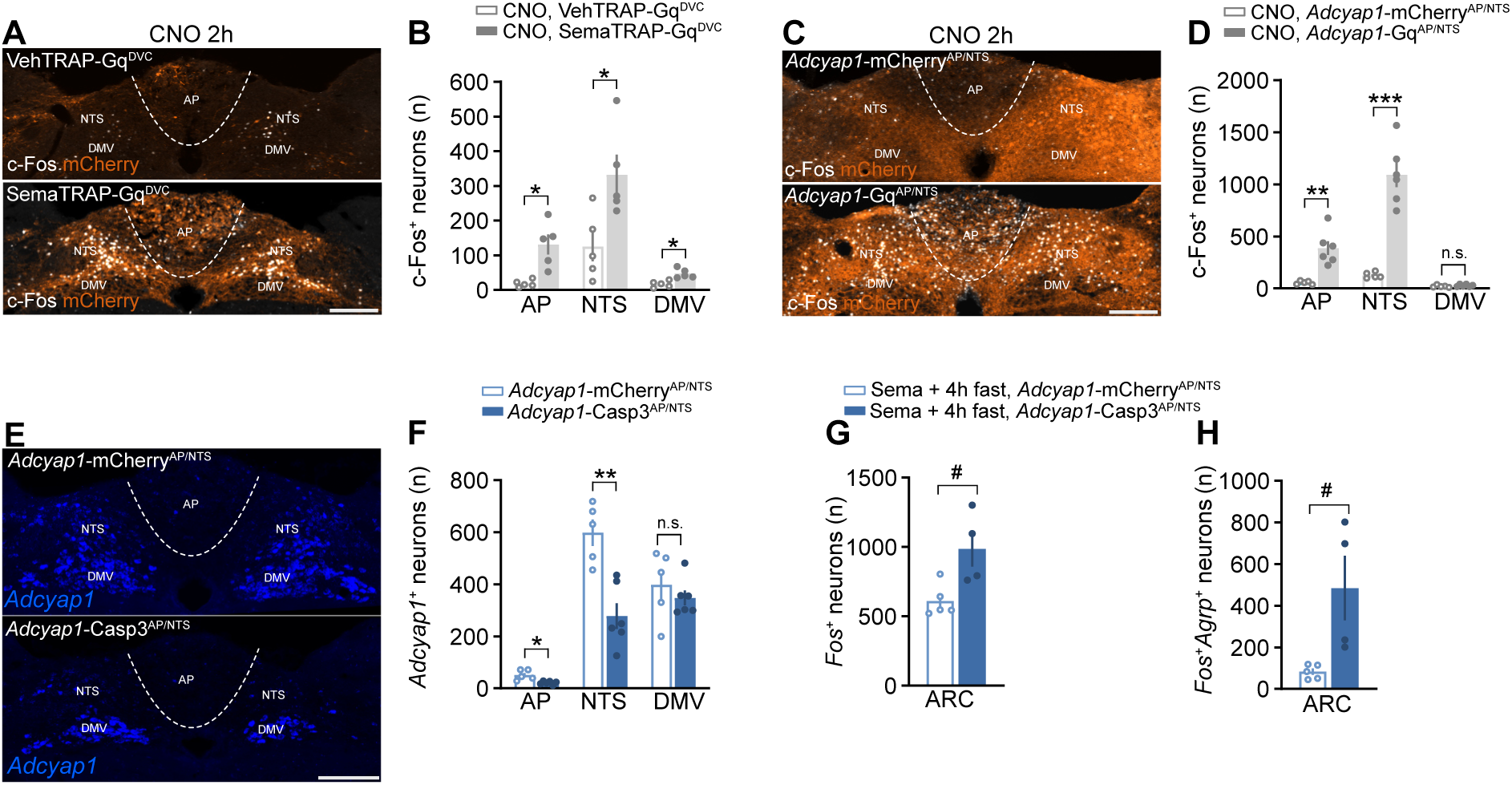
(A) AAV expression (mCherry; orange) and CNO-induced c-Fos expression (white) in the DVC of vehTRAP-Gq^DVC^ and semaTRAP-Gq^DVC^ mice. (B) Quantification of c-Fos expression in (A). *n=5*. (C) AAV expression (mCherry; orange) and CNO-induced c-Fos expression (white) in the DVC of *Adcyap1*-mCherry^AP/NTS^ and *Adcyap1*-Gq^AP/NTS^ mice. (D) Quantification of c-Fos expression in (C). *n=5-6*. (E) Fluorescent *in situ* hybridization showing *Adcyap1* mRNA expression (blue) in the DVC of *Adcyap1*-mCherry^AP/NTS^ and *Adcyap1*-Casp3^AP/NTS^ mice. (F) Quantification of *Adcyap1* mRNA expression in (E). *n=5-6*. (G, H) Quantification of *Fos* mRNA expression (G) and co-expression between *Fos* and *Agrp* mRNA (H) in the ARC of *Adcyap1*-mCherry^AP/NTS^ and *Adcyap1*-Casp3^AP/NTS^ injected with semaglutide and fasted into the darkphase. *n=4-5*. n.s.=not significant, \**p*<0.05, \*\**p*<0.01, \*\*\**p*<0.001. Data were analyzed using unpaired t-tests (G, H) or with unpaired t-tests with Holm-Šidák’s method to correct for multiple comparisons (B, D, F). Error bars represent ± SEM. Scale bars = 200 µm. AP, area postrema; cc, central canal; DMV, dorsal motor nucleus of the vagus; DVC, dorsal vagal complex; NTS, nucleus of the solitary tract. Relates to main Figure 2.

**Figure S2.**
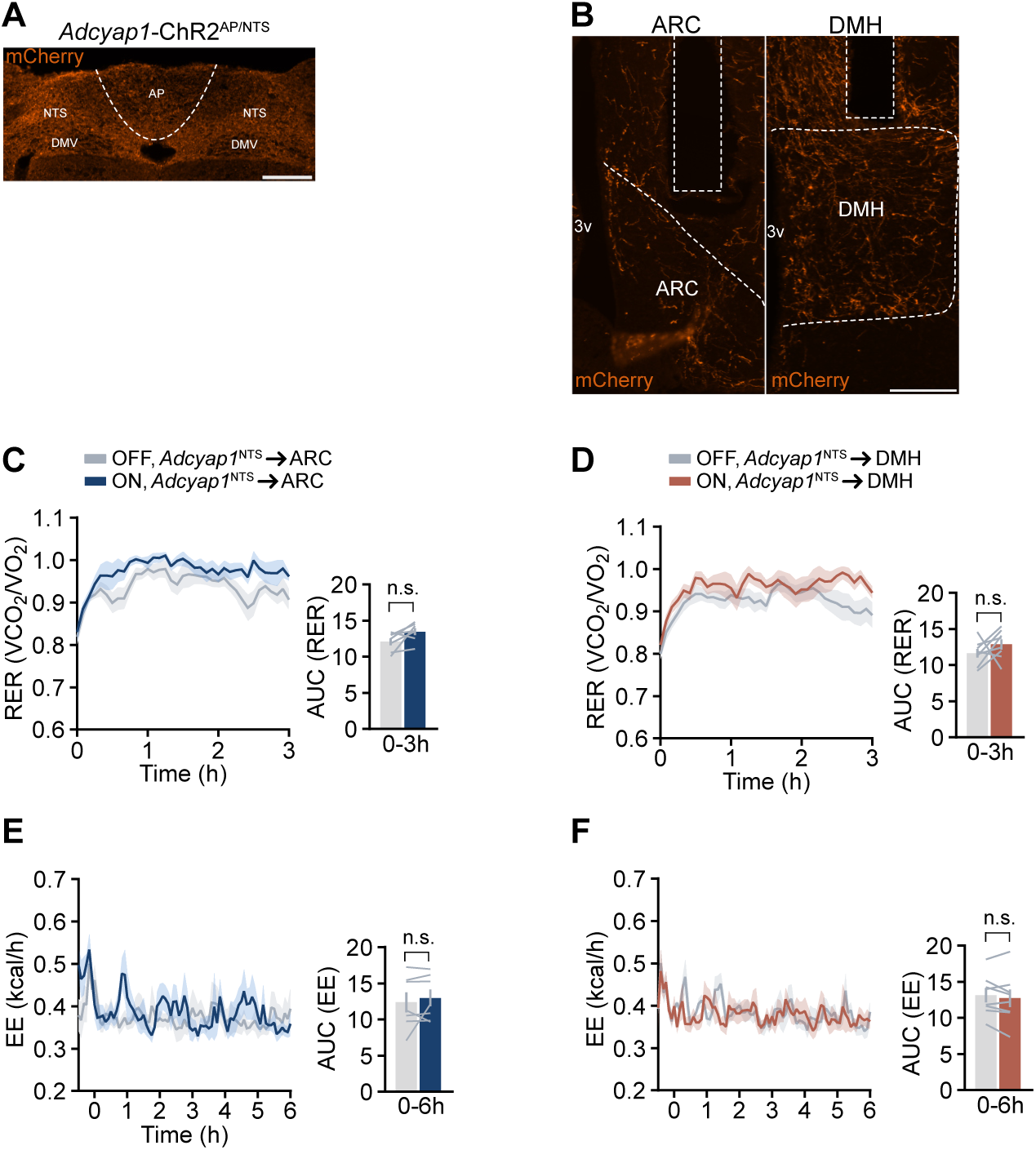
(A) Representative AAV expression in the DVC of an *Adcyap1*-2A-Cre mouse used for optogenetics (mCherry; orange). (B) *Adcyap1*-Cre-dependent expression of mCherry in NTS-derived axonal projections in the ARC and DMH, plus examples of fiberoptic cannula placements above the ARC or DMH. (C, D) Dark phase respiratory exchange ratio in photostimulated and non-stimulated *Adcyap1*^NTS^→ARC mice (C) and *Adcyap1*^NTS^→DMH mice (D). *n=7* and *n=8*. (E, F) Energy expenditure in photostimulated and non-stimulated *Adcyap1*^NTS^→ARC (E) and *Adcyap1*^NTS^→DMH mice (F) in the absence of feeding during the light phase. *n=7* and *n=8*. Data were analyzed using paired t-test (C-F). n.s.=not significant. Error bars represent ± SEM. Scale bars = 200 µm. AP, area postrema; ARC, arcuate nucleus; cc, central canal; DMH, dorsomedial hypothalamus; DMV, dorsal motor nucleus of the vagus; NTS, nucleus of the solitary tract; 3v, third ventricle. Relates to main Figure 3.

**Figure S3:**
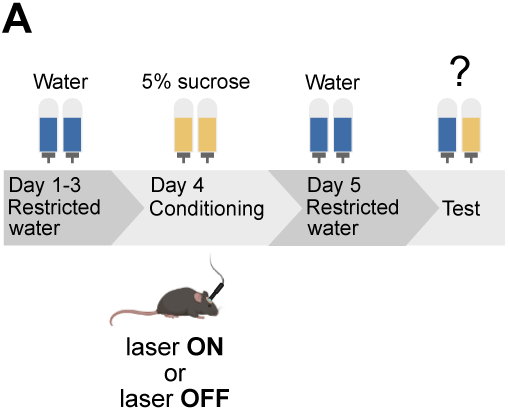
(A) Protocol used for conditioned taste aversion in combination with optogenetics. On the conditioning day, mice received photostimulation for 1 h after 30 minutes of access to a 5% sucrose solution. Controls received no laser light. Relates to main Figure 4.

**Figure S4:**
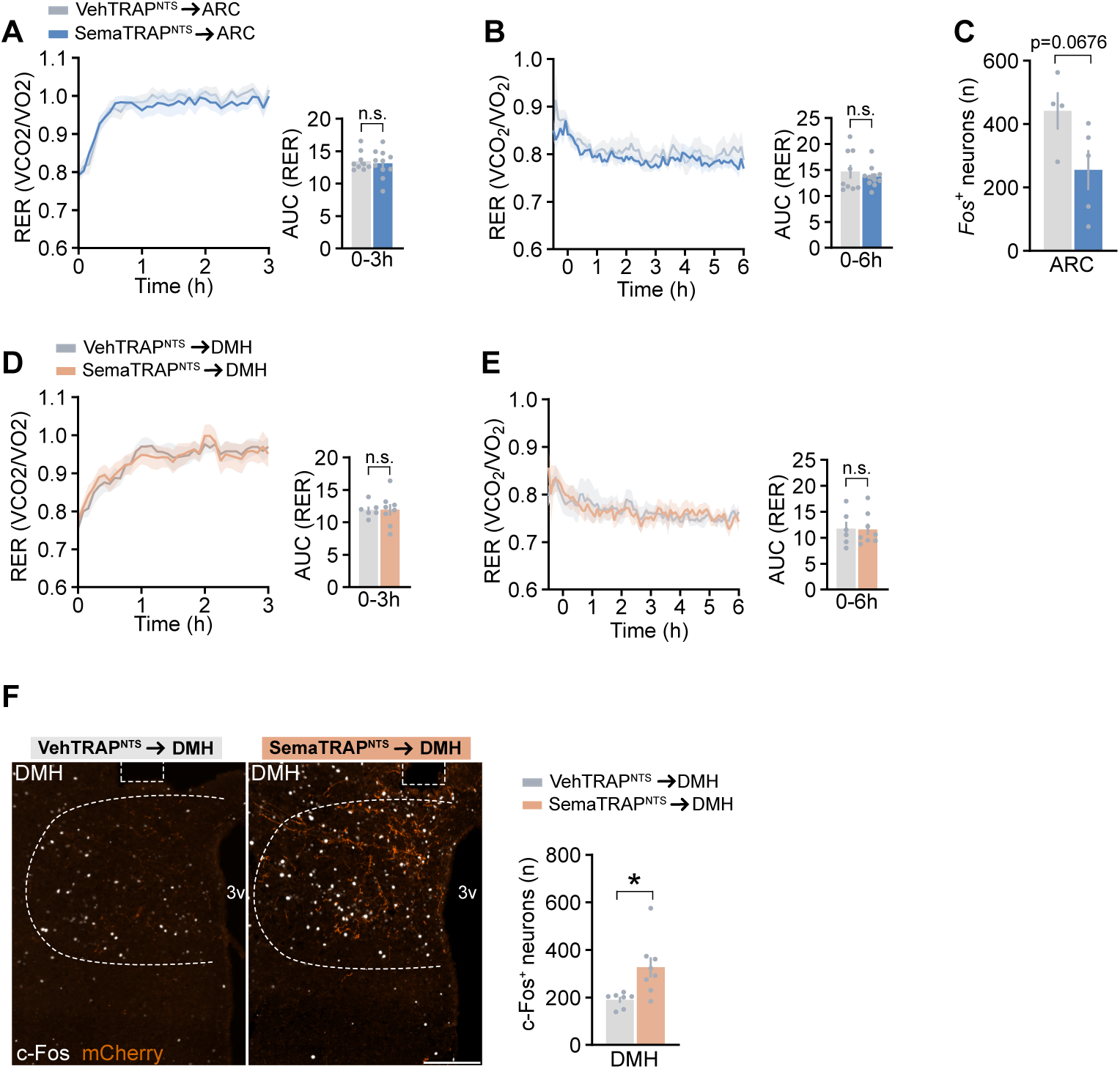
(A) Dark-phase respiratory exchange ratio during 3 h of photostimulation in vehTRAP^NTS^→ARC and semaTRAP^NTS^→ARC mice. *n=9-11*. (B) Respiratory exchange ratio during 6h of photostimulation of vehTRAP^NTS^→ARC and semaTRAP^NTS^→ARC mice in the absence of feeding during the light phase. *n=9-11*. (C) Quantification of c-Fos expressing neurons in the ARC upon photostimulation during a short fast into the dark phase in vehTRAP^NTS^→ARC and semaTRAP^NTS^→ARC mice. *n=4-5*. (D) Dark phase respiratory exchange ratio during 3 h of photostimulation in vehTRAP^NTS^→DMH and semaTRAP^NTS^→DMH mice. *n=6-8*. (E) Respiratory exchange ratio during 6h of photostimulation of vehTRAP^NTS^→DMH and semaTRAP^NTS^→DMH mice in the absence of feeding during the light phase. *n=6-8*. (F) Presence of mCherry-containing axonal fibers in the DMH (orange) derived from TRAPed NTS neurons of vehTRAP^NTS^→DMH and semaTRAP^NTS^→DMH mice, as well as DMH c-Fos expression (white) induced by photostimulation. Optic fiber placement is indicated above the DMH. Bar graph shows quantification of c-Fos expression. *n=7-8*. Scale bar = 200 µm. n.s.=not significant \**p*<0.05. Data was analyzed using unpaired t-test (A-F). ARC, arcuate nucleus; DMH, dorsomedial hypothalamus; 3v, third ventricle. Relates to main Figure 6.

## REFERENCES

1. Ryan, D.H., Lingvay, I., Deanfield, J., Kahn, S.E., Barros, E., Burguera, B., Colhoun, H.M., Cercato, C., Dicker, D., Horn, D.B., et al. (2024). Long-term weight loss effects of semaglutide in obesity without diabetes in the SELECT trial. Nat Med 30, 2049–2057. 10.1038/s41591-024-02996-7.

2. Wilding, J.P.H., Batterham, R.L., Calanna, S., Davies, M., Van Gaal, L.F., Lingvay, I., McGowan, B.M., Rosenstock, J., Tran, M.T.D., Wadden, T.A., et al. (2021). Once-Weekly Semaglutide in Adults with Overweight or Obesity. N Engl J Med 384, 989–1002. 10.1056/NEJMoa2032183.

3. Sisley, S., Gutierrez-Aguilar, R., Scott, M., D’Alessio, D.A., Sandoval, D.A., and Seeley, R.J. (2014). Neuronal GLP1R mediates liraglutide’s anorectic but not glucose-lowering effect. J Clin Invest 124, 2456–2463. 10.1172/JCI72434.

4. Imbernon, M., Saponaro, C., Helms, H.C.C., Duquenne, M., Fernandois, D., Deligia, E., Denis, R.G.P., Chao, D.H.M., Rasika, S., Staels, B., et al. (2022). Tanycytes control hypothalamic liraglutide uptake and its anti-obesity actions. Cell Metab 34, 1054–1063 e1057. 10.1016/j.cmet.2022.06.002.

5. Hansford, R., Buller, S., Tsang, A.H., Benoit, S., Roberts, A.G., Erskine, E., Brown, T., Pirro, V., Reimann, F., Harada, N., et al. (2025). Glucose-dependent insulinotropic polypeptide receptor signaling in oligodendrocytes increases the weight-loss action of GLP-1R agonism. Cell Metab 37, 1820–1834 e1825. 10.1016/j.cmet.2025.07.009.

6. Webster, A.N., Becker, J.J., Li, C., Schwalbe, D.C., Kerspern, D., Karolczak, E.O., Bundon, C.B., Onoharigho, R.A., Crook, M., Jalil, M., et al. (2024). Molecular connectomics reveals a glucagon-like peptide 1-sensitive neural circuit for satiety. Nat Metab 6, 2354–2373. 10.1038/s42255-024-01168-8.

7. Kim, K.S., Park, J.S., Hwang, E., Park, M.J., Shin, H.Y., Lee, Y.H., Kim, K.M., Gautron, L., Godschall, E., Portillo, B., et al. (2024). GLP-1 increases preingestive satiation via hypothalamic circuits in mice and humans. Science, eadj2537. 10.1126/science.adj2537.

8. Rupp, A.C., Tomlinson, A.J., Affinati, A.H., Yacawych, W.T., Duensing, A.M., True, C., Lindsley, S.R., Kirigiti, M.A., MacKenzie, A., Polex-Wolf, J., et al. (2023). Suppression of food intake by Glp1r/Lepr-coexpressing neurons prevents obesity in mouse models. J Clin Invest 133. 10.1172/JCI157515.

9. McMorrow, H.E., Cohen, A.B., Lorch, C.M., Hayes, N.W., Fleps, S.W., Frydman, J.A., Xia, J.L., Samms, R.J., and Beutler, L.R. (2025). Incretin receptor agonism rapidly inhibits AgRP neurons to suppress food intake in mice. J Clin Invest 135. 10.1172/JCI186652.

10. Huang, K.P., Acosta, A.A., Ghidewon, M.Y., McKnight, A.D., Almeida, M.S., Nyema, N.T., Hanchak, N.D., Patel, N., Gbenou, Y.S.K., Adriaenssens, A.E., et al. (2024). Dissociable hindbrain GLP1R circuits for satiety and aversion. Nature. 10.1038/s41586-024-07685-6.

11. Teixidor-Deulofeu, J., Blid Skoldheden, S., Font-Girones, F., Fejes, A., Ruud, J., and Engstrom Ruud, L. (2025). Semaglutide effects on energy balance are mediated by Adcyap1(+) neurons in the dorsal vagal complex. Cell Metab 37, 1530–1546 e1536. 10.1016/j.cmet.2025.04.018.

12. Yacawych, W.T., Wang, Y., Zhou, G., Hassan, S., Kernodle, S., Sass, F., DeVaux, M., Wu, I., Rupp, A., Tomlinson, A.J., et al. (2025). A single dorsal vagal complex circuit mediates the aversive and anorectic responses to GLP1R agonists. bioRxiv. 10.1101/2025.01.21.634167.

13. Gao, C., Geneve, I.C., Rodriguez-Gonzalez, S., Li, C., McElhern, K., Reitman, M.L., Lutas, A., and Krashes, M.J. (2026). Semaglutide drives weight loss through cAMP-dependent mechanisms in GLP1R-expressing hindbrain neurons. Nat Metab 8, 1330–1349. 10.1038/s42255-026-01534-8.

14. Cowley, M.A., Smith, R.G., Diano, S., Tschop, M., Pronchuk, N., Grove, K.L., Strasburger, C.J., Bidlingmaier, M., Esterman, M., Heiman, M.L., et al. (2003). The distribution and mechanism of action of ghrelin in the CNS demonstrates a novel hypothalamic circuit regulating energy homeostasis. Neuron 37, 649–661. 10.1016/s0896-6273(03)00063-1.

15. Steculorum, S.M., Ruud, J., Karakasilioti, I., Backes, H., Engström Ruud, L., Timper, K., Hess, M.E., Tsaousidou, E., Mauer, J., Vogt, M.C., et al. (2016). AgRP Neurons Control Systemic Insulin Sensitivity via Myostatin Expression in Brown Adipose Tissue. Cell 165, 125–138. 10.1016/j.cell.2016.02.044.

16. Douglass, A.M., Kucukdereli, H., Madara, J.C., Wang, D., Wu, C., Lowenstein, E.D., Tao, J., and Lowell, B.B. (2025). Acute and circadian feedforward regulation of agouti-related peptide hunger neurons. Cell Metab 37, 708–722 e705. 10.1016/j.cmet.2024.11.009.

17. Garfield, A.S., Shah, B.P., Burgess, C.R., Li, M.M., Li, C., Steger, J.S., Madara, J.C., Campbell, J.N., Kroeger, D., Scammell, T.E., et al. (2016). Dynamic GABAergic afferent modulation of AgRP neurons. Nat Neurosci 19, 1628–1635. 10.1038/nn.4392.

18. Brierley, D.I., Holt, M.K., Singh, A., de Araujo, A., McDougle, M., Vergara, M., Afaghani, M.H., Lee, S.J., Scott, K., Maske, C., et al. (2021). Central and peripheral GLP-1 systems independently suppress eating. Nat Metab 3, 258–273. 10.1038/s42255-021-00344-4.

19. Krashes, M.J., Koda, S., Ye, C., Rogan, S.C., Adams, A.C., Cusher, D.S., Maratos-Flier, E., Roth, B.L., and Lowell, B.B. (2011). Rapid, reversible activation of AgRP neurons drives feeding behavior in mice. J Clin Invest 121, 1424–1428. 10.1172/JCI46229.

20. Yang, C.F., Chiang, M.C., Gray, D.C., Prabhakaran, M., Alvarado, M., Juntti, S.A., Unger, E.K., Wells, J.A., and Shah, N.M. (2013). Sexually dimorphic neurons in the ventromedial hypothalamus govern mating in both sexes and aggression in males. Cell 153, 896–909. 10.1016/j.cell.2013.04.017.

21. Ilanges, A., Shiao, R., Shaked, J., Luo, J.D., Yu, X., and Friedman, J.M. (2022). Brainstem ADCYAP1(+) neurons control multiple aspects of sickness behaviour. Nature 609, 761–771. 10.1038/s41586-022-05161-7.

22. Kawai, Y. (2018). Differential Ascending Projections From the Male Rat Caudal Nucleus of the Tractus Solitarius: An Interface Between Local Microcircuits and Global Macrocircuits. Front Neuroanat 12, 63. 10.3389/fnana.2018.00063.

23. Ter Horst, G.J., de Boer, P., Luiten, P.G., and van Willigen, J.D. (1989). Ascending projections from the solitary tract nucleus to the hypothalamus. A Phaseolus vulgaris lectin tracing study in the rat. Neuroscience 31, 785–797. 10.1016/0306-4522(89)90441-7.

24. Martinez de Morentin, P.B., Gonzalez, J.A., Dowsett, G.K.C., Martynova, Y., Yeo, G.S.H., Sylantyev, S., and Heisler, L.K. (2024). A brainstem to hypothalamic arcuate nucleus GABAergic circuit drives feeding. Curr Biol 34, 1646–1656 e1644. 10.1016/j.cub.2024.02.074.

25. Tsang, A.H., Nuzzaci, D., Darwish, T., Samudrala, H., and Blouet, C. (2020). Nutrient sensing in the nucleus of the solitary tract mediates non-aversive suppression of feeding via inhibition of AgRP neurons. Mol Metab 42, 101070. 10.1016/j.molmet.2020.101070.

26. Shapiro, R.E., and Miselis, R.R. (1985). The central neural connections of the area postrema of the rat. J Comp Neurol 234, 344–364. 10.1002/cne.902340306.

27. Matikainen-Ankney, B.A., Earnest, T., Ali, M., Casey, E., Wang, J.G., Sutton, A.K., Legaria, A.A., Barclay, K.M., Murdaugh, L.B., Norris, M.R., et al. (2021). An open-source device for measuring food intake and operant behavior in rodent home-cages. Elife 10. 10.7554/eLife.66173.

28. Cavalcanti-de-Albuquerque, J.P., Bober, J., Zimmer, M.R., and Dietrich, M.O. (2019). Regulation of substrate utilization and adiposity by Agrp neurons. Nat Commun 10, 311. 10.1038/s41467-018-08239-x.

29. Gabery, S., Salinas, C.G., Paulsen, S.J., Ahnfelt-Ronne, J., Alanentalo, T., Baquero, A.F., Buckley, S.T., Farkas, E., Fekete, C., Frederiksen, K.S., et al. (2020). Semaglutide lowers body weight in rodents via distributed neural pathways. JCI Insight 5. 10.1172/jci.insight.133429.

30. Godschall, E.N., Gungul, T.B., Sajonia, I.R., Buyukaksakal, A.K., Li, O., Ogilvie, S., Keeler, A.B., Tian, G., Shi, Y., Koita, O., et al. (2026). A brain reward circuit inhibited by next-generation weight-loss drugs in mice. Nature 654, 1055–1064. 10.1038/s41586-026-10444-4.

31. Lee, S.J., Sanchez-Watts, G., Krieger, J.P., Pignalosa, A., Norell, P.N., Cortella, A., Pettersen, K.G., Vrdoljak, D., Hayes, M.R., Kanoski, S.E., et al. (2018). Loss of dorsomedial hypothalamic GLP-1 signaling reduces BAT thermogenesis and increases adiposity. Mol Metab 11, 33–46. 10.1016/j.molmet.2018.03.008.

32. Dong, Y., Carty, J., Goldstein, N., He, Z., Hwang, E., Chau, D., Wallace, B., Kabahizi, A., Lieu, L., Peng, Y., et al. (2021). Time and metabolic state-dependent effects of GLP-1R agonists on NPY/AgRP and POMC neuronal activity in vivo. Mol Metab 54, 101352. 10.1016/j.molmet.2021.101352.

33. He, Z., Gao, Y., Lieu, L., Afrin, S., Cao, J., Michael, N.J., Dong, Y., Sun, J., Guo, H., and Williams, K.W. (2019). Direct and indirect effects of liraglutide on hypothalamic POMC and NPY/AgRP neurons - Implications for energy balance and glucose control. Mol Metab 28, 120–134. 10.1016/j.molmet.2019.07.008.

34. Secher, A., Jelsing, J., Baquero, A.F., Hecksher-Sorensen, J., Cowley, M.A., Dalboge, L.S., Hansen, G., Grove, K.L., Pyke, C., Raun, K., et al. (2014). The arcuate nucleus mediates GLP-1 receptor agonist liraglutide-dependent weight loss. J Clin Invest 124, 4473–4488. 10.1172/JCI75276.

35. Jones, L.A., Cross, E., Song, Y., Claxton, P., Monaco, N., Yu, Y., Trapp, S., Adriaenssens, A., and Brierley, D.I. (2026). Semaglutide-induced satiation, nausea, and food reward suppression are mediated by GLP-1 receptors in the area postrema. bioRxiv, 2026.2008.2010.744052. 10.64898/2026.08.10.744052.

36. d’Avila, M., Cavalcanti-de-Albuquerque, J., Collado-Perez, R., Liu, Z.W., Hunter, J., White, A., Schlessinger, J., D’Agostino, G., and Horvath, T.L. (2026). AgRP neurons are required for the weight-lowering effects of GLP-1 receptor agonists in female mice. Proc Natl Acad Sci U S A 123, e2614476123. 10.1073/pnas.2614476123.

37. Sayar-Atasoy, N., Aklan, I., Yavuz, Y., Laule, C., Kim, H., Rysted, J., Alp, M.I., Davis, D., Yilmaz, B., and Atasoy, D. (2024). AgRP neurons encode circadian feeding time. Nat Neurosci 27, 102–115. 10.1038/s41593-023-01482-6.

38. Cai, J., Chen, J., Ortiz-Guzman, J., Huang, J., Arenkiel, B.R., Wang, Y., Zhang, Y., Shi, Y., Tong, Q., and Zhan, C. (2023). AgRP neurons are not indispensable for body weight maintenance in adult mice. Cell Rep 42, 112789. 10.1016/j.celrep.2023.112789.

39. Cheng, W., Gonzalez, I., Pan, W., Tsang, A.H., Adams, J., Ndoka, E., Gordian, D., Khoury, B., Roelofs, K., Evers, S.S., et al. (2020). Calcitonin Receptor Neurons in the Mouse Nucleus Tractus Solitarius Control Energy Balance via the Non-aversive Suppression of Feeding. Cell Metab 31, 301–312 e305. 10.1016/j.cmet.2019.12.012.

40. Cheng, W., Ndoka, E., Maung, J.N., Pan, W., Rupp, A.C., Rhodes, C.J., Olson, D.P., and Myers, M.G., Jr. (2021). NTS Prlh overcomes orexigenic stimuli and ameliorates dietary and genetic forms of obesity. Nat Commun 12, 5175. 10.1038/s41467-021-25525-3.

41. Chen, W., Mehlkop, O., Scharn, A., Nolte, H., Klemm, P., Henschke, S., Steuernagel, L., Sotelo-Hitschfeld, T., Kaya, E., Wunderlich, C.M., et al. (2023). Nutrient-sensing AgRP neurons relay control of liver autophagy during energy deprivation. Cell Metab 35, 786–806 e713. 10.1016/j.cmet.2023.03.019.

42. Takahashi, T.M., Sunagawa, G.A., Soya, S., Abe, M., Sakurai, K., Ishikawa, K., Yanagisawa, M., Hama, H., Hasegawa, E., Miyawaki, A., et al. (2020). A discrete neuronal circuit induces a hibernation-like state in rodents. Nature 583, 109–114. 10.1038/s41586-020-2163-6.

43. Pinol, R.A., Zahler, S.H., Li, C., Saha, A., Tan, B.K., Skop, V., Gavrilova, O., Xiao, C., Krashes, M.J., and Reitman, M.L. (2018). Brs3 neurons in the mouse dorsomedial hypothalamus regulate body temperature, energy expenditure, and heart rate, but not food intake. Nat Neurosci 21, 1530–1540. 10.1038/s41593-018-0249-3.

44. Duensing, A.M., Belmont-Rausch, D., Tomlinson, A.J., Crowley, A., Sass, F., Heaton, E., Coester, B., Brown, J.M., Hassan, S., Wu, Z., et al. (2026). Molecularly defined subpopulations of leptin receptor neurons dissociate the control of food intake from blood pressure. bioRxiv. 10.64898/2026.03.26.714551.

45. Tong, Q., Ye, C.P., Jones, J.E., Elmquist, J.K., and Lowell, B.B. (2008). Synaptic release of GABA by AgRP neurons is required for normal regulation of energy balance. Nat Neurosci 11, 998–1000. 10.1038/nn.2167.

46. Allen, W.E., DeNardo, L.A., Chen, M.Z., Liu, C.D., Loh, K.M., Fenno, L.E., Ramakrishnan, C., Deisseroth, K., and Luo, L. (2017). Thirst-associated preoptic neurons encode an aversive motivational drive. Science 357, 1149–1155. 10.1126/science.aan6747.

47. Zhu, H., Aryal, D.K., Olsen, R.H., Urban, D.J., Swearingen, A., Forbes, S., Roth, B.L., and Hochgeschwender, U. (2016). Cre-dependent DREADD (Designer Receptors Exclusively Activated by Designer Drugs) mice. Genesis 54, 439–446. 10.1002/dvg.22949.

48. Harris, J.A., Hirokawa, K.E., Sorensen, S.A., Gu, H., Mills, M., Ng, L.L., Bohn, P., Mortrud, M., Ouellette, B., Kidney, J., et al. (2014). Anatomical characterization of Cre driver mice for neural circuit mapping and manipulation. Front Neural Circuits 8, 76. 10.3389/fncir.2014.00076.

49. Ye, L., Allen, W.E., Thompson, K.R., Tian, Q., Hsueh, B., Ramakrishnan, C., Wang, A.C., Jennings, J.H., Adhikari, A., Halpern, C.H., et al. (2016). Wiring and Molecular Features of Prefrontal Ensembles Representing Distinct Experiences. Cell 165, 1776–1788. 10.1016/j.cell.2016.05.010.

50. Chen, J.Y., Campos, C.A., Jarvie, B.C., and Palmiter, R.D. (2018). Parabrachial CGRP Neurons Establish and Sustain Aversive Taste Memories. Neuron 100, 891–899 e895. 10.1016/j.neuron.2018.09.032.

51. Bankhead, P., Loughrey, M.B., Fernandez, J.A., Dombrowski, Y., McArt, D.G., Dunne, P.D., McQuaid, S., Gray, R.T., Murray, L.J., Coleman, H.G., et al. (2017). QuPath: Open source software for digital pathology image analysis. Sci Rep 7, 16878. 10.1038/s41598-017-17204-5.

